# BCL6 regulates the endothelial pro-immunogenic phenotype relevant to organ transplant rejection

**DOI:** 10.1101/2022.11.03.514941

**Authors:** Adriana Franco Acevedo, Nicole M Valenzuela

**Affiliations:** Department of Pathology and Laboratory Medicine, University of California, Los Angeles, USA

## Abstract

**Background:** IFNγ induces an endothelial cell pro-immunogenic phenotype through the JAK/STAT1 pathway, which can influence alloreactive leukocytes in transplant rejection. Numerous endogenous suppressors of JAK and STAT activation have been described, but regulation of STAT1 at the level of transcription is not well-understood. In immune cells, the DNA binding protein BCL6 controls transcription of lineage and inflammatory genes, including STAT-dependent responses. The goal of this study was to determine if BCL6 also modulates the IFNγ-induced immunogenic phenotype in endothelium.

**Methods:** BCL6 binding to target genes and similarity to STAT1 motifs was analyzed *in silico* using public datasets. *In vitro*, primary human aortic endothelial cells were tested for expression of IFNγ-inducible costimulatory molecules, HLA and cytokines, under BCL6 overexpression, depletion and pharmacological targeting. Gene expression was measured by RNA-Seq and protein expression was confirmed by flow cytometry, Luminex and ELISA. Paired biopsies from stable and rejecting human cardiac allografts were compared for expression of BCL6, PD ligands, CXCL chemokines and HLA-DR in the vasculature.

**Results:** BCL6 expression is increased within human cardiac transplants during rejection, which is positively correlated with expression of interferon response gene HLA-DR in donor blood vessels. Further, BCL6 is IFNγ-inducible via JAK1/2 in endothelium. Next, the consensus DNA binding motif of BCL6 is highly similar to that of STAT1, and numerous interferon response genes harbor BCL6 DNA binding motifs. Depletion of BCL6 in endothelium results in augmentation, while overexpression causes suppression, of MHC class II, chemokine, and PD ligand expression. Unexpectedly, pharmacological targeting of the corepressor domain BTB domain of BCL6 also repressed many interferon response genes, particularly HLA class II, CXCR3 chemokines and PD-L2. On the other hand, BCL6 targeting did not impact inducible expression of any HLA class I genes, PD-L1 or CD40; and the effect is correlated with the presence of BCL6 binding motifs in or near affected genes.

**Conclusion:** Our results show for the first time that the transcriptional repressor BCL6 selectively controls the endothelial response to IFNγ. A better understanding of the endogenous mechanisms that regulate donor endothelial activation has the potential to discover new avenues to dampen transplant rejection, with broader relevance to other vascular inflammatory diseases.

## Introduction

Transplantation of heart, kidney, lung and liver allografts is a lifesaving treatment for end-stage organ failure. However, recipients must be maintained on lifelong immunosuppression to prevent rejection of the transplanted organ. The primary target of the rejection response is the donor endothelial cell, which is directly exposed to the recipient’s immune system. Endothelial cells are critical for maintenance of vascular integrity and regulation leukocyte trafficking into peripheral tissues ^6^. Additionally, endothelial cells are increasingly recognized as semi-professional antigen presenting cells, capable of expressing a limited repertoire of costimulatory molecules, cytokines and T cell receptor ligands that can activate and skew adaptive immune cells, including T cells ^1–5^.

Genetic mismatches in the major histocompatibility complex between donor and recipient result in potent alloimmune recognition of foreign antigen by the recipient adaptive immune system, including both T cell and B cells/antibodies. In humans, classical HLA class I molecules are encoded by HLA-A, -B, and –C genes, as well as minor histocompatibility antigens MICA and MICB. HLA class I molecules are constitutively expressed by all nucleated cells, including donor vascular cells, and can be further upregulated under several inflammatory conditions, including after exposure to TNFα or IFNγ. HLA class II molecules include HLA-DR, -DQ and -DP, and accessory molecules CD74, DO, and DM; but these are expressed only by professional antigen presenting cells, and conditionally by endothelium. In particular, HLA-DQ mismatches, and donor specific antibodies to this protein, confer the highest risk of rejection and graft loss. In addition to HLA, endothelial cells can express a modest repertoire of costimulatory and coinhibitory molecules, such as CD40, PD ligands, and cytokines, to influence T cell activation ^1–5^. The balance of these molecules is an important determinant in alloimmune activation or tolerance.

Although it exerts complex and contradictory effects on allograft outcome depending on the compartment and context, animal models have demonstrated that IFNγ production by infiltrating leukocytes ^6, 7^ and donor IFNγ responsiveness contribute to acute and chronic rejection ^8, 9^. Transplant rejection can be reduced by deficiency of IFNγ-stimulated effectors such as HLA class II or CXCL10 within the transplanted organ ^10, 11^. On the other hand, intragraft PD ligands attenuate alloimmune responses ^12, 13^. Deletion of IFNγ receptors in the graft results in little to no expression of MHC and low immune cell infiltration ^14^. IFNγ activates the transcription factors STAT1 and IRF1 to trigger expression of interferon stimulated genes (ISGs). Reflecting the importance of this pathway, antagonism of intragraft STAT1 ameliorates rejection of heart transplants in mice ^15^. Despite the importance to transplant rejection, endogenous direct repressors of interferon-induced transcription and mediators of chromatin remodeling remain to be identified.

Transcriptome analysis of clinical transplant biopsies revealed a strong IFNγ-dependent mRNA signature within grafts ^14, 16–18^. Interferons predominantly activate JAK/STAT pathways, with IFNγ acting through JAK1 and JAK2 to promote transcription via IRF1 and STAT1. Mechanistically, IFNγ induces several costimulatory molecules and HLA II in endothelium, to promote transition to an antigen presentation phenotype ^4, 9, 10^. We previously demonstrated that this response is durable or self-sustaining from prime-withdrawal experiments ^4^, and is exclusively dependent on JAK1/2 signaling. Despite the importance of donor endothelial expression of HLA and costimulatory molecules in transplantation, knowledge is lacking on transcriptional modulators of the endothelial semi-antigen presenting phenotype.

We identified that the gene *BCL6* is differentially expressed in endothelial cells from various anatomic origins, and is particularly enriched in cardiac artery endothelium ^19^. BCL6 is a widely expressed DNA binding protein that regulates transcription of target genes, through recognition of target sequences by its zinc finger domains. Its function is mainly characterized in the adaptive immune compartment, where it is a lineage-defining factor for T follicular helper cells (Tfh) and involved in germinal center formation ^20, 21^. Oncogenic mutations in BCL6 are common and causative in B cell lymphoma ^22^. Further, BCL6 controls expression of inflammatory mediators in macrophages ^23, 24^ and metabolic processes in adipocytes ^25^]. One major mechanism of action, at least in the immune compartment, is association with corepressors and histone deacetylases to form a transcriptional repressor complex, via protein-protein interactions through the BTB/POZ domain. Nevertheless, loss or perturbation of BCL6 results in both activation/derepression and suppression of hundreds of genes in numerous cell types.

BCL6 is IFNγ-inducible via STAT1 in several cell types ^26–31^. BCL6 is also known to regulate STAT-dependent responses in B cells, and to modulate antiviral signaling/IFN ^32^, and regulate PD ligand expression in B cells ^33^. However, very little is known about the function of BCL6 in vascular cells. It appears to control a subset of VEGF-induced genes during angiogenesis ^34, 35^. Additionally, BCL6 was shown to bind to the *VCAM1* gene and to suppress expression of other inflammatory genes ^36, 37^. However, despite its importance in regulating inflammatory and immune responses in innate and adaptive hematopoietic cells, nearly nothing is known about the role of BCL6 in regulating endothelial inflammatory responses.

Based on this knowledge, we hypothesized that BCL6 specifically regulates STAT1-dependent IFNγ transcriptional programming in endothelium. We found that BCL6 is highly increased in human cardiac allograft biopsies undergoing rejection in comparison with stable allografts, and correlated with expression of interferon response gene HLA-DR. In human cardiac endothelial cells, inducible antigen presentation and costimulatory molecule expression were tested under overexpression, knockdown or pharmacological targeting of BCL6. These *in vitro* experiments demonstrate a previously unreported role for BCL6 in repressing IFNγ-induced endothelial gene expression, particularly MHC class II, CXCR3 chemokines and PD ligands. The results show that BCL6 is a significant regulator of the endothelial pro-immunogenic phenotype. Surprisingly, antagonism of the BTB domain of BCL6 enhanced its repressive effects similar to BCL6 overexpression. Thus, BCL6 targeting is a novel therapeutic approach to specifically dampen allograft immunogenicity, that has the potential to prevent acute and chronic transplant rejection.

## Materials and Methods

### Ethics Statement

Use of human endothelial cells and peripheral blood mononuclear cells was approved by the UCLA IRB (IRB#17-00477). Studies involving human heart allograft samples were approved under the UCLA IRB#21-001330.

### Human cardiac allograft biopsies

Normal human tonsil and archived remnant human endomyocardial biopsy tissue from heart transplant recipients were obtained from the UCLA TPCL. Paired remnant FFPE biopsies from five patients each were selected with the following criteria: and a normal biopsy less than 3 months post-transplant and biopsy-proven acute rejection ≥ACR1R and/or ≥pAMR1 after the first 3 months post-transplant. Biopsies were stained by H&E and immunofluorescence OPAL kit for CD31, BCL6, HLA-DR, and CD45 (Dako Agilent) by the UCLA TPCL, and digitally scanned on Aperio. Normal human tonsil stained strongly positive for CD45, with diffuse HLA-DR and punctate BCL6 expression in germinal centers (**Supplemental Figure 1A**), demonstrating the specificity of the antibodies.

### Cells

Peripheral blood human mononuclear cells were purchased from PromoCell. Primary human aortic endothelial cells (HAEC) were purchased from commercial sources: PromoCell, Lonza, SciCell and ATCC. Endothelial cells were cultured in complete medium (PromoCell MV2) on tissue culture treated vessels pre-coated with 0.2% gelatin (Sigma), and used for experiments at passages 2-8. Experiments were repeated with biological replicates of primary human aortic endothelium (n≥3 donors). Reporter IFNγ (GAS) and ISG (IRF) HEK293 cell lines were purchased from Invivogen and expanded in DMEM supplemented with 10% heat inactivated FBS and selection antibiotics per vendor instructions.

### Reagents

Human IFNγ was purchased from Sigma Millipore. 79-6, FX1, BI-3812, BI-3802, and ruxolitinib were obtained from Selleck Chemical. FX1 was pre-tested at the concentration range 1-100μM, as the reported efficacy in B cells, reactivating gene expression was 50μM ^38^.

### Lentiviral BCL6 shRNA and Overexpression

We performed a pilot screen of lentiviral vectors which identified hCMV and mCMV as the most active promoters in human aortic endothelial cells (Promoter Screen, Horizon Dharmacon) (**Supplemental Figure 5**). For knockdown, cells were transduced with pLenti BCL6 shRNA GFP or negative control GAPDH shRNA (Horizon Dharmacon). For overexpression, cells were transduced with pLenti BCL6 ORF-mGFP or empty mGFP only (Origene).

Endothelial cells were seeded at 40-60% confluence, incubated overnight, and transduced with third generation lentiviral plasmids encoding LentiORF-BCL6 (mGFP-tagged), LentiORF control (mGFP-P2A-Puro) (Origene), shRNA against BCL6 or GAPDH shRNA by incubating with lentiviral particles at 25-100 MOI in complete medium with polybrene at 8μg/mL overnight. After an additional 24hr rest period in complete medium, cells were used for experiments; or, once confluent, GFP expression was confirmed, and transduced cells were expanded and used for future experiments.

### RNA Sequencing

HAEC were cultured to confluence in 6 well plates in complete medium. Where indicated, cells were pre-treated with inhibitors diluted in M199 + 10% heat inactivated FBS for 30min, followed by addition of IFNγ (200U/mL) for 24hr. Then, conditioned medium was removed, monolayers were washed once with PBS without Ca^2+^ or Mg^2+^, and detached with Accutase (vendor). Cells were pelleted at 350 xg for 6 min, and lysed in 350μL of RLT Buffer (Qiagen). After vortexing and centrifugation at 14,000 RPM at 4⁰C, lysates were stored at −20⁰C. Total RNA was purified using the RNeasy kit (Qiagen), and submitted for sequencing at the UCLA TCGB.

The reads were mapped by STAR 2.7.9a and read counts per gene were quantified using the human genome GRCh38. In Partek Flow, read counts were normalized by CPM +1.0E-4. All results of differential gene expression analysis utilized the statistical analysis tool, DESeq2. P-value (p<0.01), FDR (<0.01), and fold change (FC>2-fold) filters were applied for differentially expressed gene lists prior to downstream analysis. Using the list of significantly differentially expressed genes, the Canonical Pathway analysis, Disease & Function analysis, and Networks analysis were performed in QIAGEN Ingenuity Pathway Analysis software (IPA).

### Immunofluorescence Microscopy

Endothelial cells were cultured to confluence in gelatin-coated, black well/glass bottom, tissue culture treated vessels (ibidi), then stimulated as indicated. Medium was removed, cells were washed once with PBS, and then fixed with 4% paraformaldehyde for 15min at room temperature. After fixation, cells were permeabilized with 0.01% in PBS, blocked with goat serum and stained with rabbit anti-BCL6 (Cell Signaling, 1:100) and mouse anti-STAT1 (Santa Cruz), followed by anti-rabbit-AF647 and anti-mouse-AF488 (Jackson Immuno, 1:500). Images were acquired at 4X on a Cytation5 multimodal plate reader.

### Western Blotting

Endothelial cells were cultured to confluence in 6 well tissue culture treated plates and stimulated as indicated. Medium was removed, cells were washed once with PBS with 1X Halt Protease Inhibitor, and 200μL of Lysis Buffer (Cell Signaling Technologies) supplemented with 1X Halt Protease Inhibitor (ThermoFisher) was added to adherent cells on ice. Cells were detached and disrupted by scraping, and lysates were clarified by centrifugation at 14,000 x*g* at 4⁰C for 10min.

After addition of Laemmli buffer, lysates were boiled, separated on a Criterion TGX gel (Biorad), and transferred to PVDF membrane. Membranes were blotted with rabbit anti-BCL6 (Cell Signaling Technologies).

### Flow cytometry: Endothelial Cells

HAEC were cultured to confluence in 24 or 48 well plates in complete medium. Where indicated, cells were pre-treated with inhibitors diluted in M199 + 10% heat inactivated FBS for 30min, followed by addition of IFNγ (200U/mL) for 24hr. Then, conditioned medium was removed, monolayers were washed once with PBS without Ca^2+^ or Mg^2+^, and detached with Accutase (vendor). Cells were pelleted at 350 xg for 6 min, resuspended in antibody mix in FACS buffer (PBS + 2% hi-FBS), and stained for 45min at 4⁰C in the dark. After washing once, fluorescence was measured on a Fortessa flow cytometer (BD Biosciences). Data were analyzed in FlowJo.

Panel 1: HLA-ABC-BV510, HLA-DR-BV421, PD-L1-FITC or CD40-FITC, PD-L2-APC, BST2-PE/Cy7. Panel 2: HLA-ABC-BV510, HLA-DR-BV421, HLA-DQ1,3,4-FITC, HLA-DQ2,3-FITC, HLA-DP-PE, PD-L1-PE/Cy7, PD-L2-APC (Biolegend, BD Biosciences). For transduced cells, FITC/AF488 markers were omitted due to expression of GFP.

### Flow Cytometry: PBMC

PBMC were cultured in RPMI + 20% FBS alone or with IFNγ (200U/mL) for 24hr and 48hr. Cells were resuspended, including a wash with PBS to collect adherent monocytes, and stained for subset markers (CD3-APC/Fire, CD19-APC, CD14-PerCP/Cy5.5, CD56-PE/Cy7) and HLA (HLA-ABC-BV510, HLA-DR-BV421, HLA-DQ1,3,4-FITC, HLA-DQ2,3-FITC, and HLA-DP-PE (Biolegend, BD Biosciences) for 45min at 4⁰C in the dark. After washing, fluorescence was measured on a Fortessa flow cytometer (BD Biosciences). Data were analyzed in FlowJo by gating on lymphocytes and monocytes using FSC/SSC, followed by CD3+CD19-T cells, CD3-CD19+ B cells, CD3-CD19-CD14+ monocytes, and CD3-CD19-CD14-CD56+ NK cells. Then, the median fluorescence of cell surface HLA proteins was determined for each gated subset.

### Protein Phosphorylation

HAEC monolayers were cultured in 6 well plates and treated as above. After washing with PBS, cells were lysed directly in the plates in 350μL of ice-cold cell lysis buffer. Lysates were pre-cleared by high speed centrifugation at 4⁰C. Phosphorylation of STAT proteins was assayed by Luminex (Milliplex STAT) at the UCLA Immune Assessment Core.

### Quantification of Secreted Chemokines

Conditioned medium was stored at −20⁰C, then assayed in technical duplicate for secreted chemokines and cytokines by ELISA (IL-15, BAFF, CXCL10/IP-10, and CXCL11/I-TAC, all R&D Systems; CXCL9/MIG, Invitrogen) according to the manufacturer’s protocol. Additionally, supernatants were tested for 38 chemokines and cytokines by Luminex (Milliplex) at the UCLA Immune Assessment Core.

### GAS/ISG Reporter Cell Lines

HEK293 stably transfected with an SEAP reporter (HEK-Dual and HEK-Blue) responsive to STAT1 (IFNγ/GAS) or IRF (ISG) were obtained from Invivogen. Cells were seeded in a 96 well plate and treated with IFNγ alone or in the presence of BCL6 inhibitors. Transcription factor activity was read out with SEAP substrate according to the vendor’s instructions.

### Public Dataset Analysis

We analyzed two public microarray datasets of interferon-treated endothelial cells: GSE106524, GSE3920, ^39, 40^. Data were accessed in NCBI GEO and analyzed in GEO2R. BCL6 ChIP-Seq data in HEK293 and HepG2 cells were analyzed in ENCODE ^41^. We used ChIP Atlas Enrichment Analysis ^42^ to perform an unbiased query of transcription factors binding to consensus DNA motifs of interest. Specifically, we analyzed TF binding to the BCL6, STAT1 and IRF1 degenerate sequences from MotifMap ^43^, with the following parameters: q <1E-05 vs random permutation (x1) of same motif.

### Statistical Analyses

Technical replicates are defined as repeated measures of the same analyte with the same biological specimen. Biological replicates are defined as repeated measures of the same analyte with biological specimens from different sources and/or different experiments. Experiments were repeated with a minimum of three biological replicates, as indicated in the figure legends. Outliers were not excluded from analyses except where negative or positive control conditions failed. Differences between groups were compared by one way ANOVA followed by multiple comparisons testing using GraphPad Prism. Heat maps and hierarchical clustering were generated using Morpheus (https://software.broadinstitute.org/morpheus/).

## Results

### BCL6 expression in the rejecting human heart transplant

We recently reported that *BCL6* was among 27 immune-regulatory genes that were enriched in cardiac endothelial cells ^19^. To understand its physiological role, we investigated whether BCL6 expression correlated with vascular immunogenicity during transplant rejection. To that end, paired endomyocardial biopsies from heart transplant recipients were analyzed, comparing a stable biopsy and a biopsy with acute rejection from each of 5 patients. Patient demographics and biopsy diagnoses can be found in **Table 1.**

**Table 1.**

| Case | Pre-formed DSA | Biopsy post-operative day | Diagnosis | DSA at Bx |
| --- | --- | --- | --- | --- |
| 1 | A2, MICA | 59<br>930 | No rejection<br>No rejection | None<br>dnDQ2 4 mo prior |
| 2 | None | 482<br>1252 | No rejection<br>No rejection | Weak dnDSA DQ8<br>None for 2yr prior |
| 3 | None | 62<br>395<br>423 | No rejection<br>ACR2R, pAMR<br>indeterminate<br>pAMR1 h+ | None<br>dnDSA strong DQ<br>Persistent strong DQ DSA |
| 4 | A68, B51 | 34<br>247<br>1717 | pAMR1 i+<br>pAMR2<br>pAMR1 h+ | Persistent A/B, dnDSA DQ<br>Persistent A DSA<br>Strong >1 dnDSA since 2.5yr prior |
| 5 | None | 58<br>699<br>905 | No rejection<br>ACR1R<br>pAMR2 | None<br>dnDSA B53, DQ2<br>Persistent DQ2 DSA |

We stained biopsies using clinically validated antibodies against CD45, PECAM1, HLA-DR, and BCL6. Positive control tissue human tonsil highlights the specificity of the BCL6 antibody, where germinal centers stained diffusely as expected (**Supplemental Figure 1A**). Interestingly, one of the endomyocardial biopsies contained a Quilty lesion (intragraft tertiary lymphoid organ), noted by the diagnosing pathologist as CD19+ B cell rich. We found that these lesions contained numerous cells strongly positive for CD45 and BCL6, further supporting the specificity of the antibody staining (**Supplemental Figure 1B, 1C**).

Exemplary H&E and immunofluorescence micrographs from each case are shown in **Figure 1**. Over all 5 cases we observed the same pattern, where BCL6, HLA-DR and CD45 were rarely detected in endomyocardial biopsies not diagnosed with rejection (**Figure 1E-1J**)). In contrast, numerous CD45+ infiltrates were seen in rejection biopsies. Further, expression of HLA-DR was enhanced in acute rejection biopsies, including colocalizing with CD31+ blood vessels (**Figure H**). Notably, diffuse BCL6 positivity was also apparent in all biopsies with rejection, again, with some staining localizing to inflamed CD31+ endothelium (**Figure 1H**). As an example, Case 5 had a normal biopsy early post-transplant, but one year post-transplant, developed de novo DQ DSA and graft dysfunction. Biopsy on POD382 showed ACR2R, and biopsy on POD423 had residual AMR1h+ (**Figure 1D**). Very little CD45, BCL6 or HLA-DR was detected in the stable biopsy taken POD62 without rejection, but extensive HLA-DR and BCL6 staining was found in the myocardium in the biopsy with pAMR2 (**Figure 1J**).

**Figure 1.**
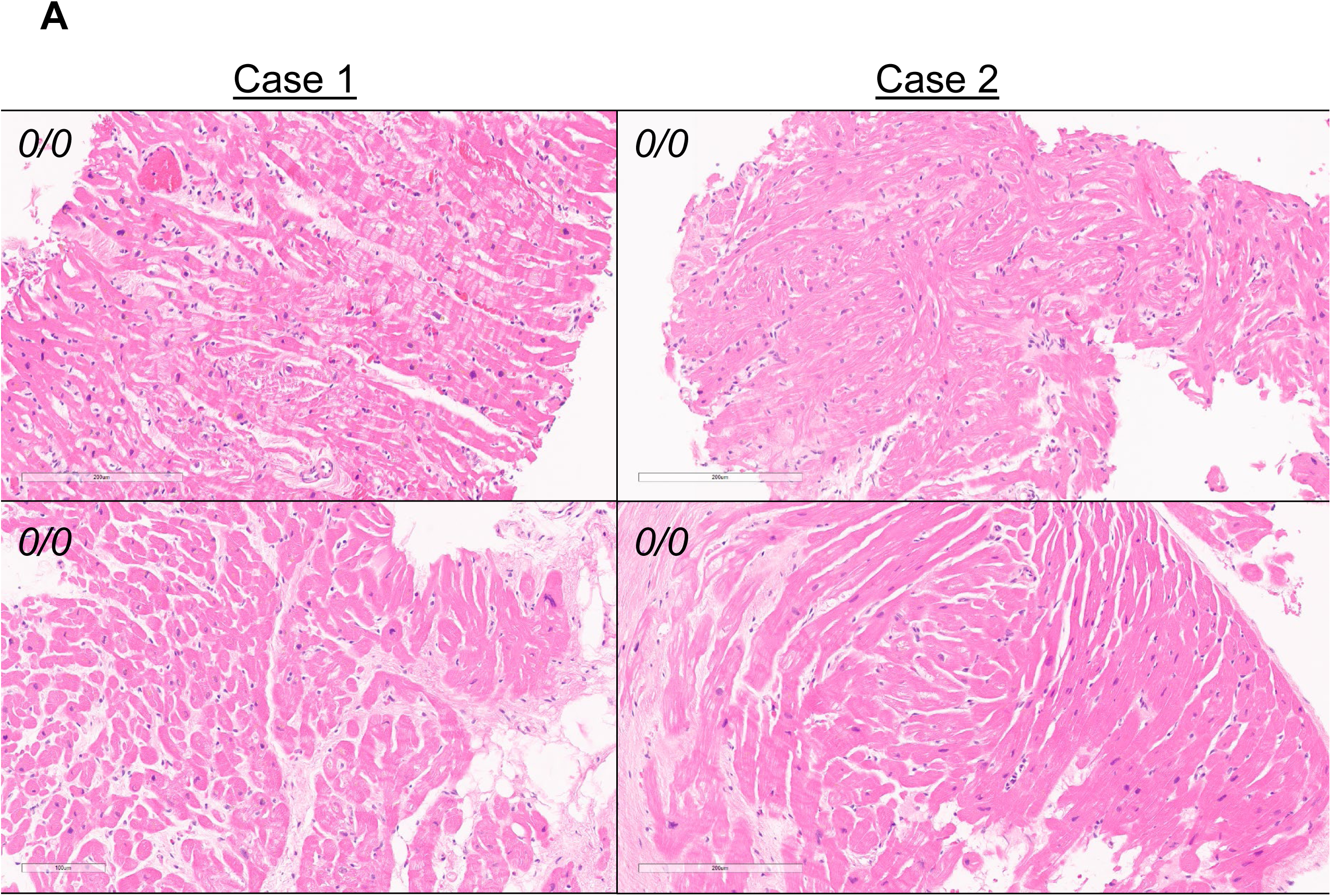
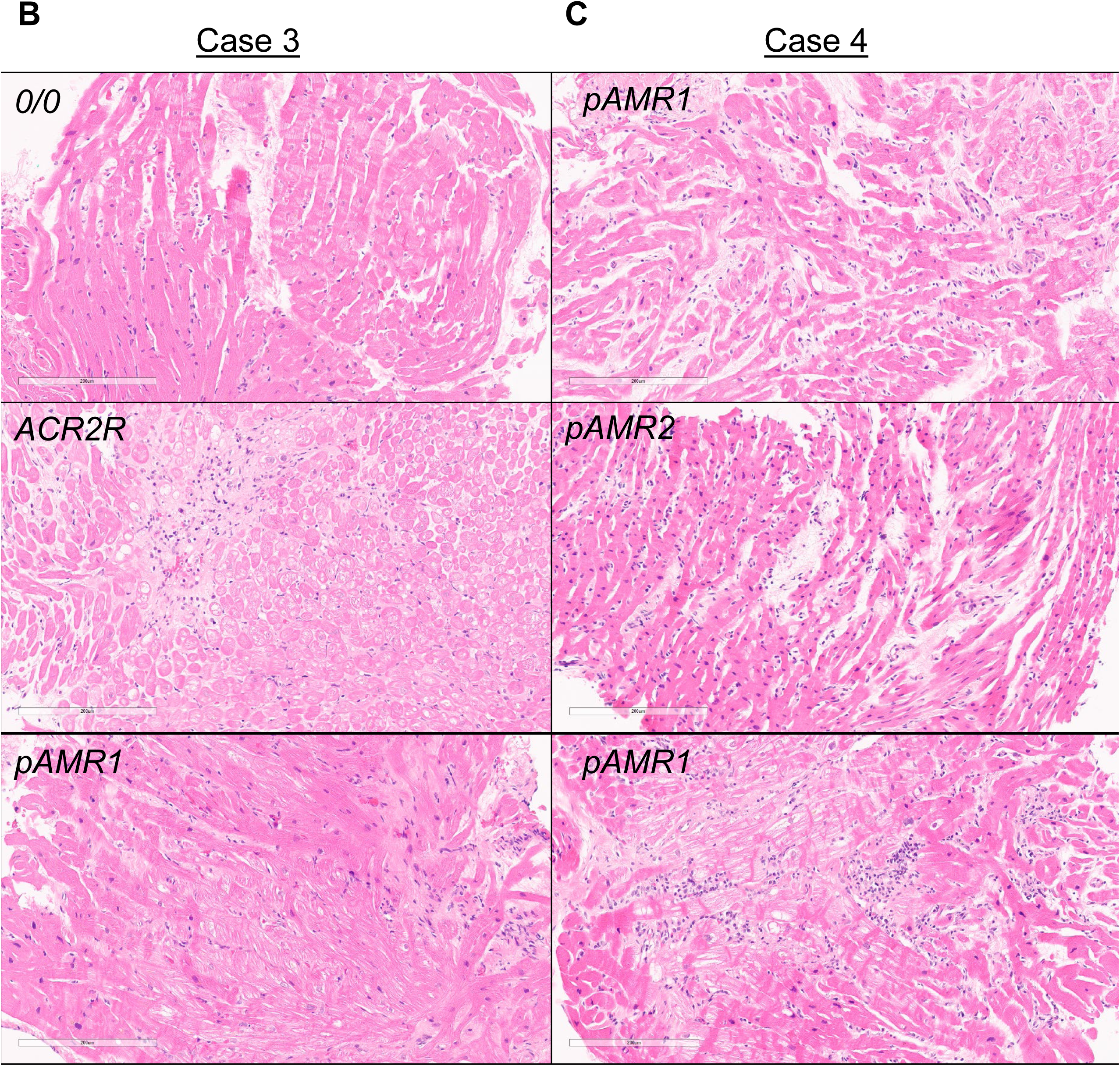
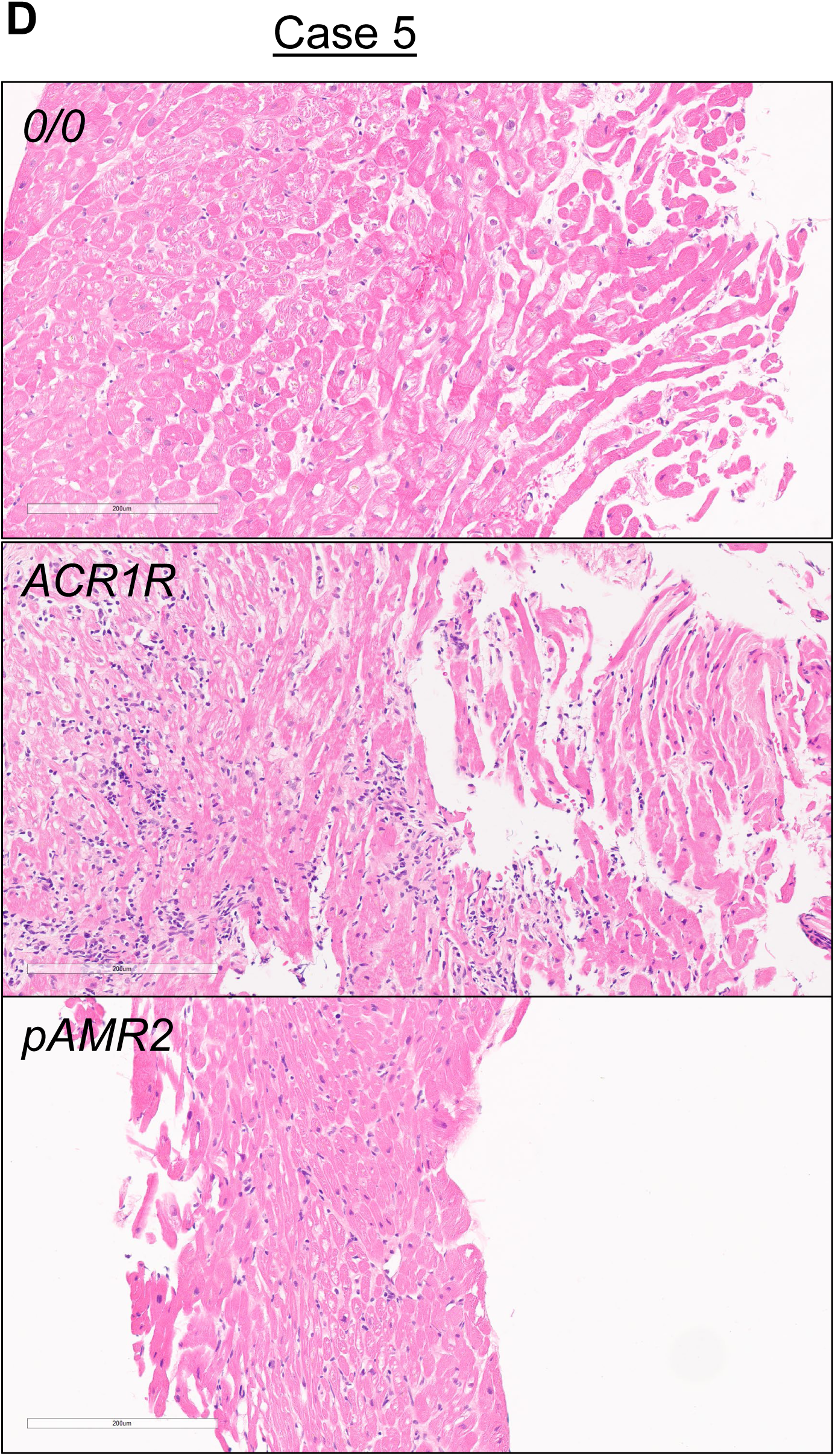
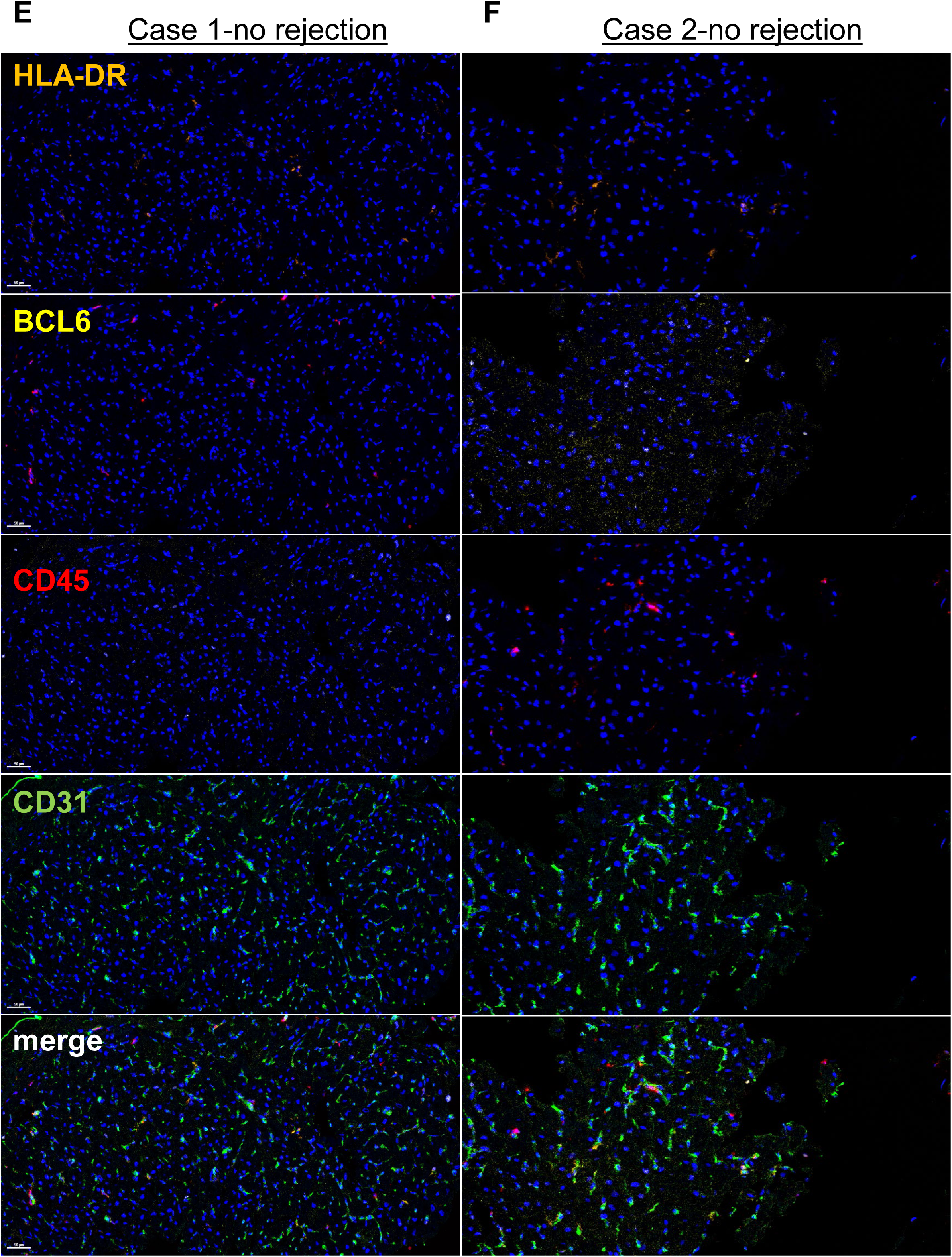
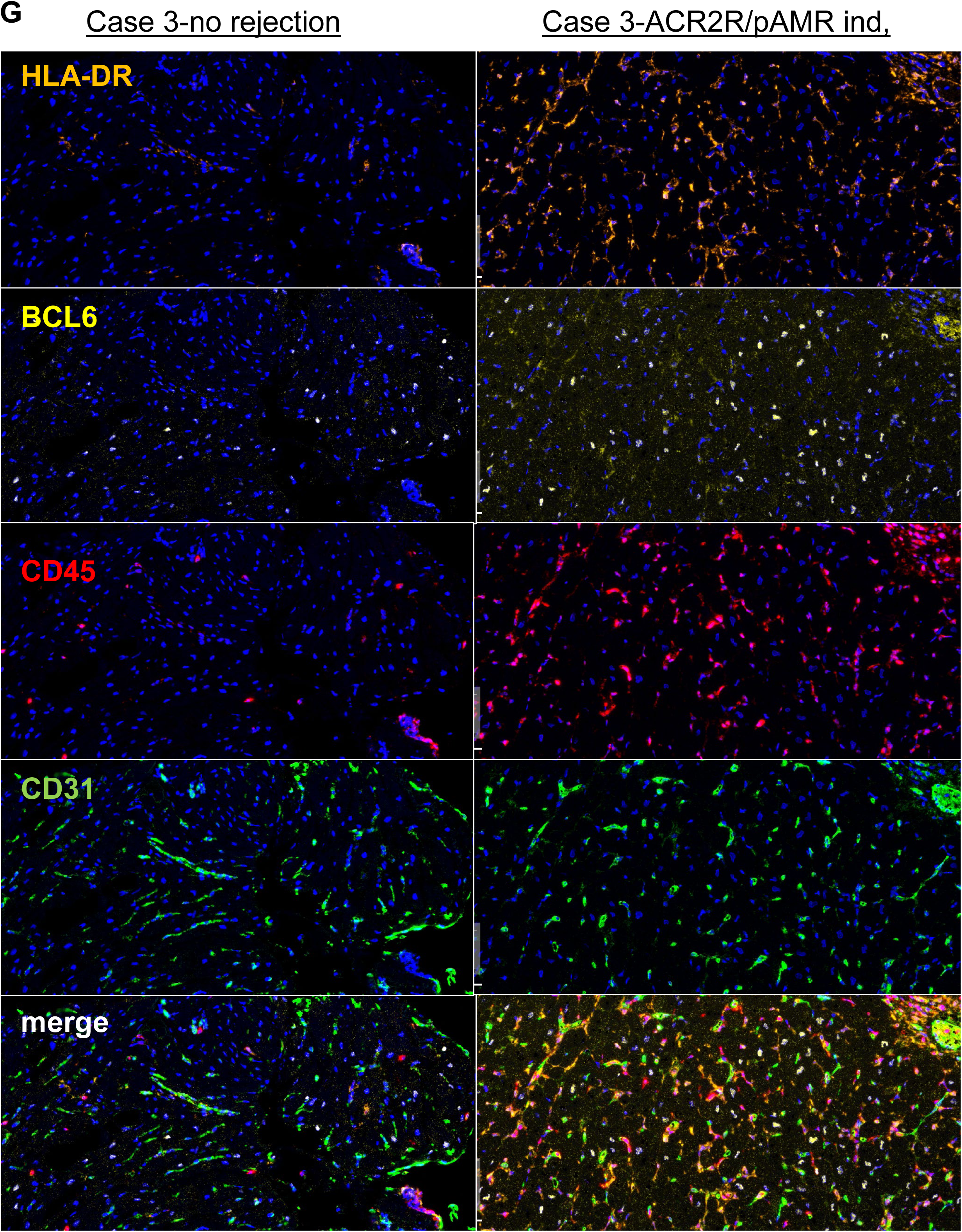
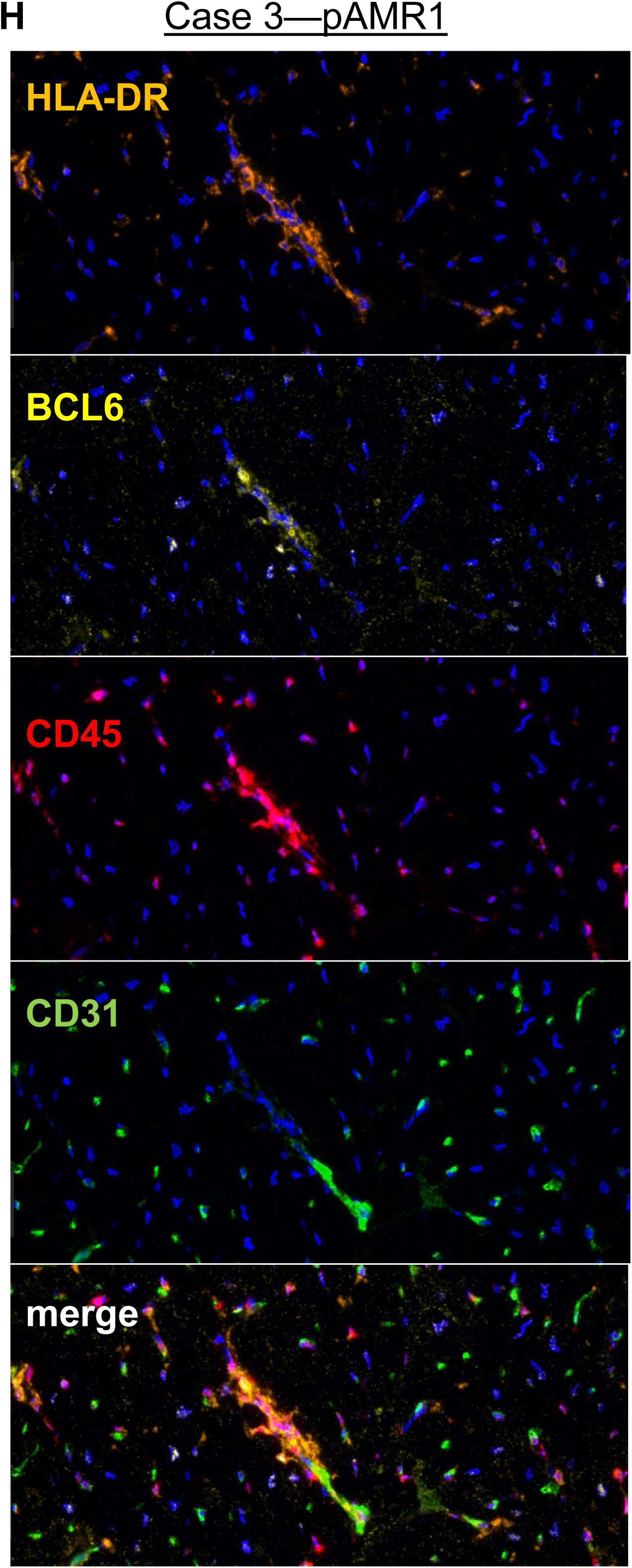
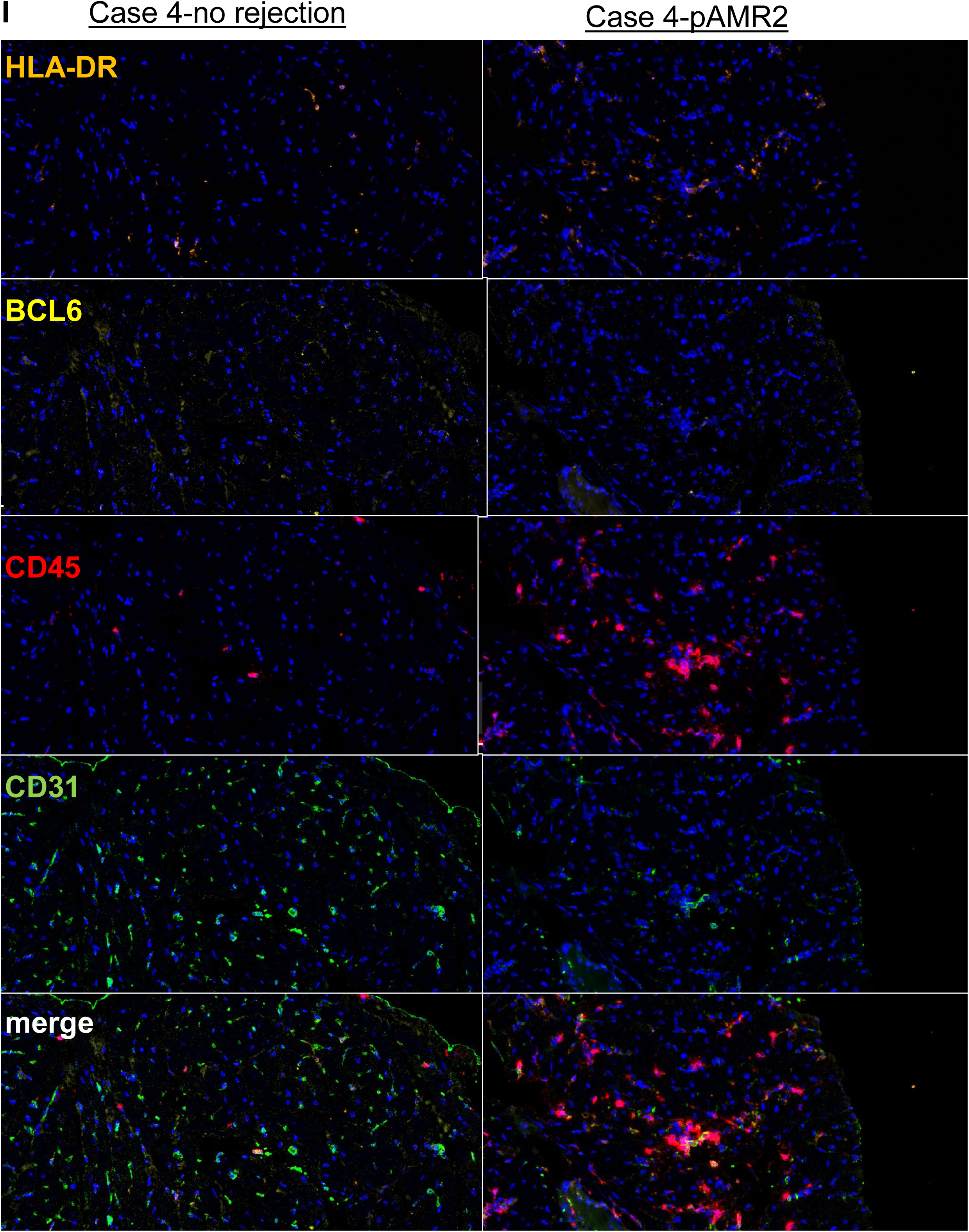
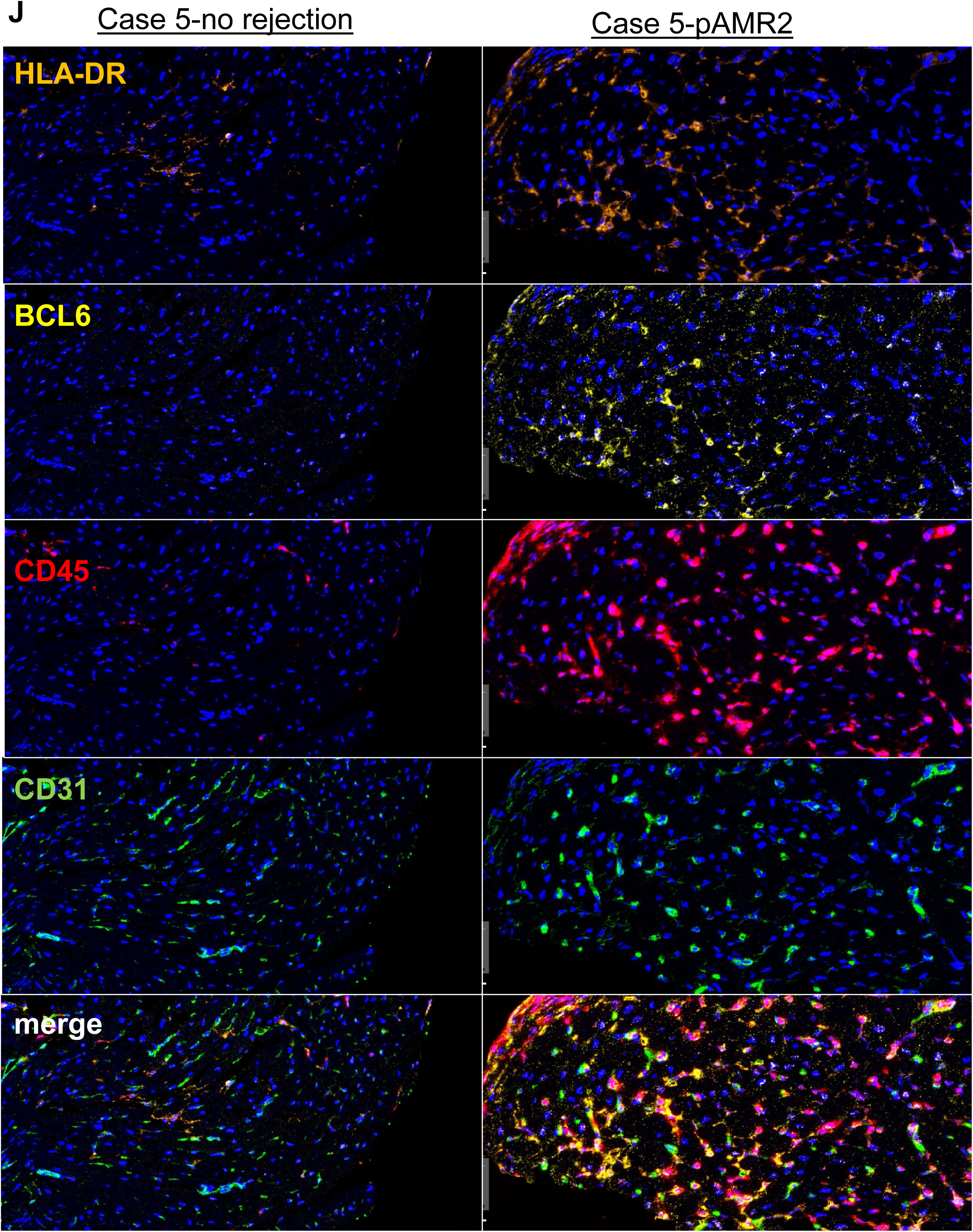
BCL6 expression is increased in human heart transplants during biopsy-proven rejection. A) Representative 20X H&E micrographs from Case #1 (POD 59, top left; POD 930, bottom left) and Case #2 (POD 482, top right; POD 1252, bottom right). Paired longitudinal biopsies from each patient are shown, each diagnosed with no rejection. B) Representative 20X H&E micrographs from Case #3. Early post-transplant protocol biopsy on POD 62 showed no rejection (top), followed by ACR2R and indeterminate diagnosis of AMR on POD 395 (middle), and follow-up resolved ACR but residual pAMR1 h+ on POD 423 (bottom). C) For Case #4, early biopsy on POD 34 diagnosed with pAMR1 i+ (top). Later the patient experienced pAMR2 on POD 247 (middle), and on POD 1717 was again diagnosed with pAMR1 h+ (bottom) D) Case #5 had a clean biopsy on POD 58 (top), but on POD 699 experienced mild rejection ACR1R (middle), and later on POD 905 was diagnosed with pAMR2 (bottom) **E-J**) Remnant FFPE endomyocardial biopsies were stained for HLA-DR (orange), BCL6 (yellow), CD45 (red), CD31 (green) and DAPI (blue) by multiplex immunofluorescence. Representative 10X micrographs are shown, except for H) which provides a 40X magnification of an inflamed vessel.

### BCL6 is IFNγ inducible in EC

Because we found that BCL6 is constitutively expressed at a low level in cardiac endothelial cells ^19^, but increased during heart transplant rejection, we asked whether BCL6 could be upregulated by inflammatory cytokines. In particular, IFNγ is a major driver of transplant rejection, and BCL6 is known to be IFNγ inducible in T follicular helper cells ^29^.

Indeed, we found that IFNγ increased BCL6 expression 2.36±0.37-fold in HAEC from 3 different donors (**Figure 2A**). Further, BCL6 transcripts were increased by IFNγ in human endothelial cells from multiple different vascular beds (HUVEC, HLMVEC, HPAEC, HRGEC and HLSEC) (**Figure 2B**). Many other ZBTB family genes were expressed at baseline in endothelial cells, some very highly (**Supplemental Figure 2**). However, none were changed by IFNγ (**Supplemental Figure 2**); only BCL6 was significantly increased.

**Figure 2.**
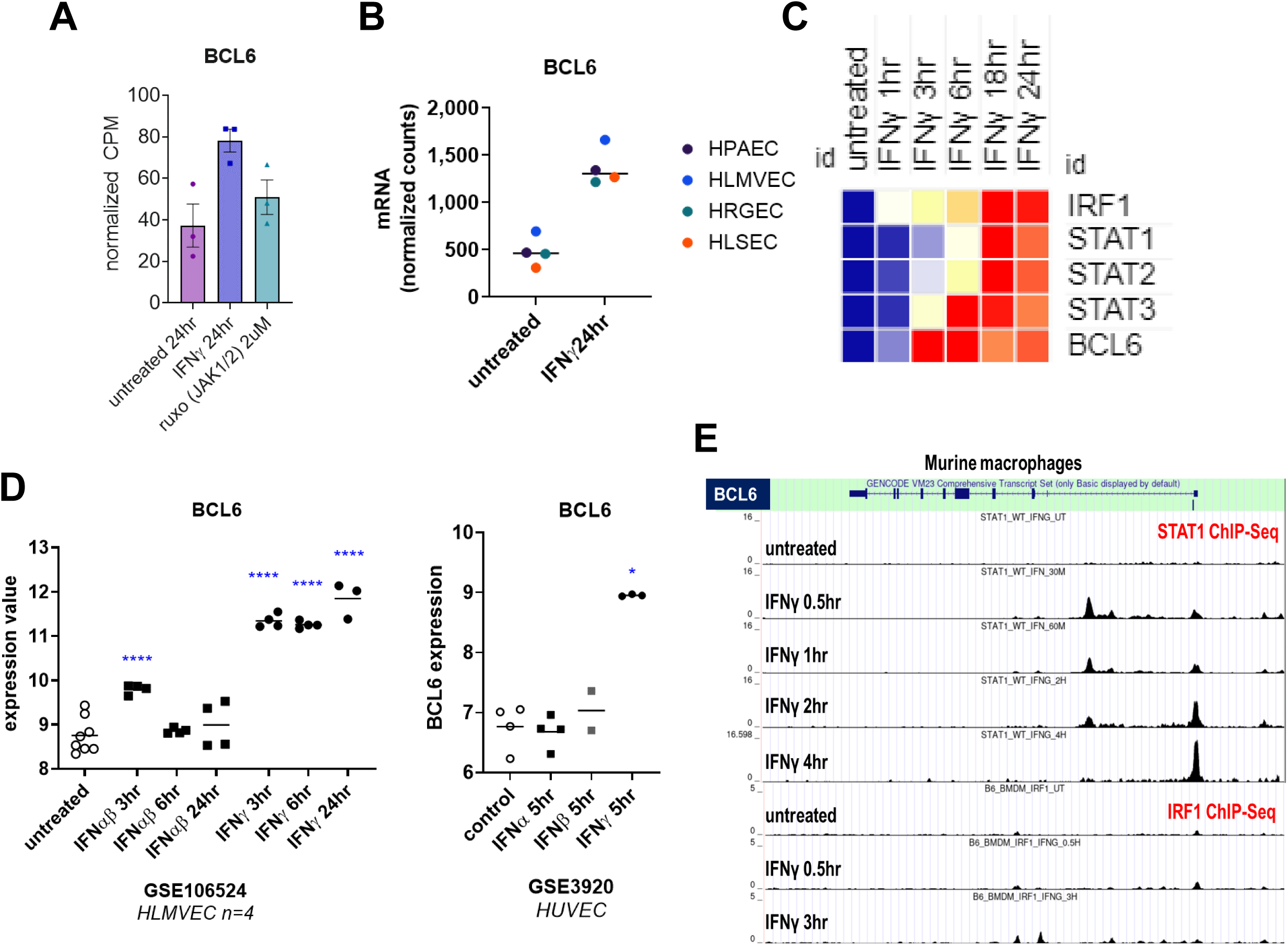
BCL6 is IFNγ inducible in EC in a JAK1/2 dependent manner. A) Human aortic endothelial cells (n=3 biological replicates) were pre-treated with ruxolitinib (2μM), then treated with IFNγ for 24hr and gene expression was measured by RNA-Seq. Normalized *BCL6* transcript CPM is graphed. B) Primary human pulmonary artery (HPAEC), lung microvascular (HLMVEC), renal glomerular (HRGEC), or liver sinusoidal (HLSEC) endothelial cells were treated with IFNγ for 24hr and gene expression was measured by Nanostring. Normalized transcript counts of *BCL6* are shown. C) Human HMEC-1 endothelial cells were treated with IFNγ for 1-24hr and gene expression was measured by Nanostring. Relative transcript counts are shown in the heat map over time. D) Left panel: Public microarray data from GSE106524 of HLMVEC (n=4) treated with IFNγ or IFNα/β for 3-24hr was analyzed in GEO2R. Right panel: Public microarray data from GSE3920 of HUVEC N=3) treated with IFNγ, IFNα or IFNβ for 5hr was analyzed in GEO2R. The mRNA expression value for *BCL6* is graphed. Groups were compared by one way ANOVA followed by multiple comparisons testing to the untreated condition. E) Public ChIP-Seq data from GSE?? of STAT1 and IRF1 transcription factor binding in murine macrophages treated with IFNγ for 0.5-3hr was visualized in the UCSC Genome Browser. ChIP peaks for each transcription factor are shown at the *Bcl6* gene.

We confirmed these results in two public datasets of human endothelial cells treated with type I or type II IFNγ [GSE106524; GSE3920]. Interestingly, human lung microvascular endothelial cells stimulated with type I IFNα/β only transiently and lowly increased BCL6 mRNA, which was not observed at 6hr or 24hr (**Figure 2D, left panel**). Similar results were found in HUVEC treated with IFNα, IFNβ, or IFNγ for 5hr, where only IFNγ-treated cells upregulated BCL6 (**Figure 2D, right panel**).

Because IFNγ but not type I IFN triggered a sustained increase in BCL6 expression, we asked whether intracellular signaling particular to type II IFN was required for BCL6 elevation. IFNγ acts through JAK1/2 to activate the transcription factors IRF1 and STAT1 homodimer, while type I IFNs use JAK1/TYK2 to activate a STAT1/STAT2 heterodimer. We first examined a public dataset of IRF1 and STAT1 genomic binding in macrophages treated with IFNγ ^44^. Here, STAT1 bound at the transcriptional start site and within exon 1 of the *Bcl6* gene within 30min of IFNγ exposure (**Figure 2E**). In contrast, no IRF1 peaks were found at any time point (**Figure 2E**). Then, we tested pre-treatment of HAEC with the JAK1/2 inhibitor ruxolitinib (2μM, 30min), and the increase in BCL6 was abrogated by ruxolitinib (**Figure 2A**). These results demonstrate that IFNγ elevates BCL6 expression in endothelium through the JAK1/2-STAT1 pathway.

### IFNγ stimulated genes in endothelial cells

We previously reported the temporal induction of IFNγ-regulated costimulatory molecules in endothelial cells ^4^. Here, we performed RNA-Seq to get a broader profile, and found that 227 genes were decreased and 475 genes were increased in primary human aortic endothelial cells by IFNγ treatment at 24hr. We treated primary human aortic endothelial cells (HAEC, n=3) with IFNγ for 24hr and compared transcriptome changes to untreated cells by RNA-Seq. IFNγ stimulation of endothelial cells significantly increased 305 genes and decreased 40 genes at 24hr (n=3 biological replicates, p<0.05, log2FC ≥2.0 or ≤-2.0). As expected, IPA analysis highlighted activation of key pathways including “antigen presentation pathway” and “interferon signaling”. Top significantly differentially expressed genes included CXCR3 chemokines, adhesion molecules and costimulatory molecules (**Figure 3A**).

**Figure 3.**
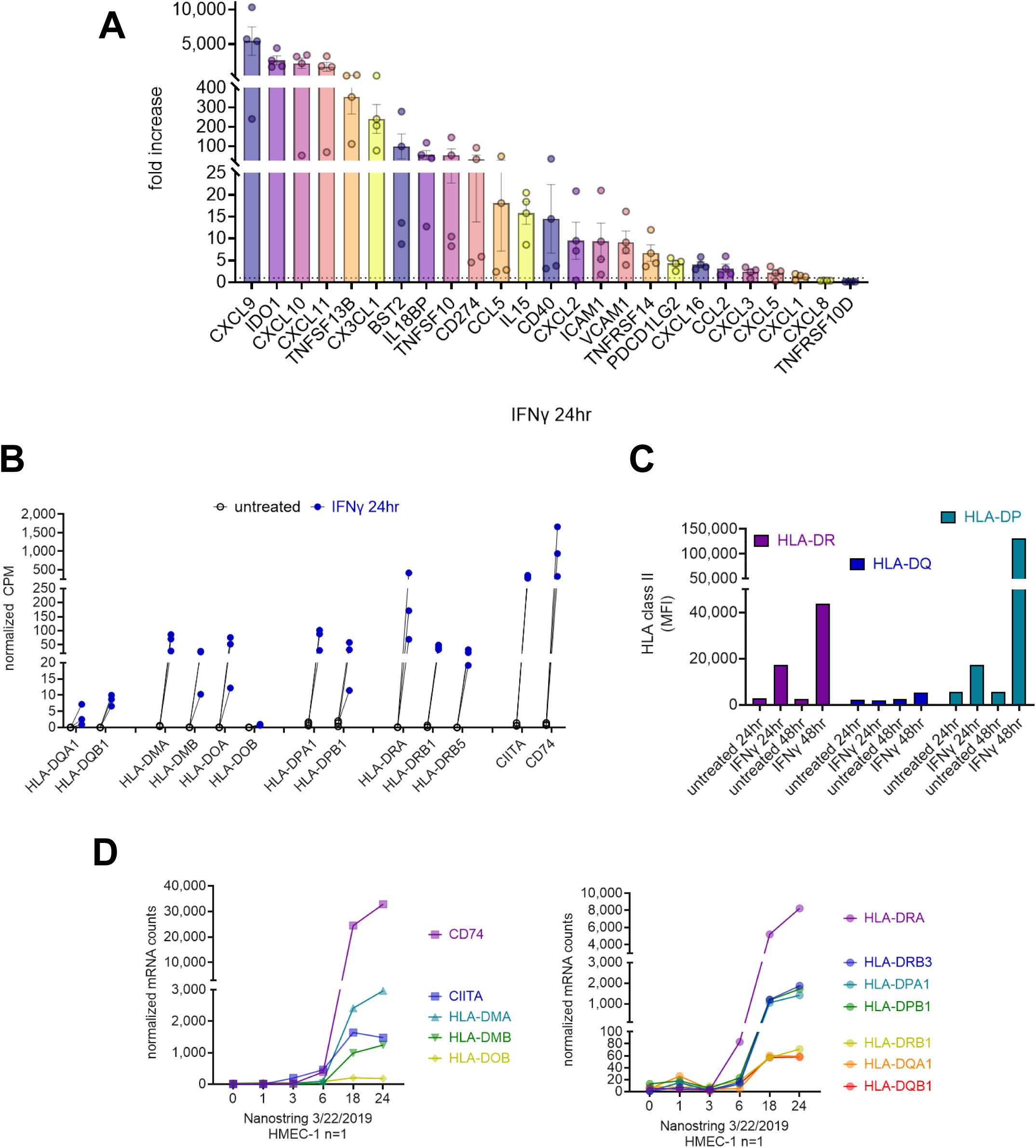
IFNγ responsive gene expression in endothelial cells is predominated by HLA, CXCR3 chemokines, and PD ligands. A) Human aortic endothelial cells (n=3 biological replicates) were treated with IFNγ for 24hr and gene expression was measured by RNA-Seq. The fold change in normalized CPM for each gene compared with untreated controls is shown. B) Normalized CPM transcript counts for HLA genes are shown, with spaghetti plots graphing the increase compared with paired untreated controls. C) Primary human aortic endothelial cells were treated with IFNγ for 24hr and 48hr. Cell surface HLA-DR, -DQ, and –DP were measured by flow cytometry. D) HMEC-1 human endothelial cells were treated with IFNγ for 1-24hr. RNA expression was measured by Nanostring. Normalized mRNA counts are graphed over time.

We noted a difference in HLA class II locus expression, where transcripts for HLA-DRA and HLA-DPB1 at 24hr were 4 to 20-fold higher than for HLA-DQA1 and DQB1 genes (**Figure 3B**). Increased expression of HLA-ABC and new expression of HLA-DR and HLA-DP proteins was detected at 24hr after IFNγ treatment; yet, no HLA-DQ was detected (**Figure 3C**). HLA-ABC, HLA-DR and HLA-DP were substantially increased at 48hr, and at this later time point, low levels of cell surface HLA-DQ protein could be detected (**Figure 3C**). The mRNA tempo (in HMEC-1, Nanostring) further identified an upregulation of HLA-DRA as early as 6hr of IFNγ treatment, and its transcript was the highest at 24hr compared with other loci. In contrast, HLA-DQA1 and HLA-DQB1 transcripts were 25 times lower than for HLA-DPA1, HLA-DPB1 and HLA-DRB3 at the same time point (**Figure 3D**).

As comparators, we tested cell surface protein HLA-DR, DQ and DP expression on PBMC from three healthy donors, left untreated or stimulated with IFNγ for 24hr and 48hr (**Supplemental Figure 3**). Untreated CD14^high^ monocytes expressed high levels of HLA class I and HLA-DR after 24hr and 48hr of culture which was similar to that on B cells, but lower levels of HLA-DQ and HLA-DP compared with untreated B cells. IFNγ slightly increased HLA-DR expression at 24hr but not further at 48hr on monocytes. Also surprising was that HLA class I was not increased by IFNγ at either time point. In contrast, monocyte expression of HLA-DQ and HLA-DP was substantially increased (-fold) after 24hr of IFNγ and further increased at 48hr. Similar results were seen with B cells, on which HLA class I and HLA-DR was not substantially affected by IFNγ, while HLA-DQ and HLA-DP were increased at 24hr and 48hr. T cells did not express appreciable levels of HLA-DR, DQ or DP in any condition. These results point to a cell type-specific regulation of HLA expression by IFNγ.

### BCL6 motif overlap with STAT1

Because BCL6 is a transcriptional repressor, we asked whether BCL6 and the major IFNγ-stimulated transcription factors, STAT1 and IRF1, might bind shared gene targets, which would suggest a role for BCL6 as a feedback regulator of IFNγ signaling. JASPAR matrix clustering showed that BCL6 motifs are very closely related to core motifs recognized by the STAT family of transcription factors ^45^. Alignment of the motifs shows substantial potential overlap (**Figure 4A**). Next, we used STAMP ^46^ to assess the similarity between the BCL6 consensus DNA binding sequence (MA0463) and STAT1. This revealed a high degree of overlap with that of the core recognition sequence of STAT (GAS motif), in both mouse and human (**Supplemental Figure 4A**). Of the top 10 most similar consensus sequences, 7 were recognized by STAT transcription factors, including STAT1.

**Figure 4.**
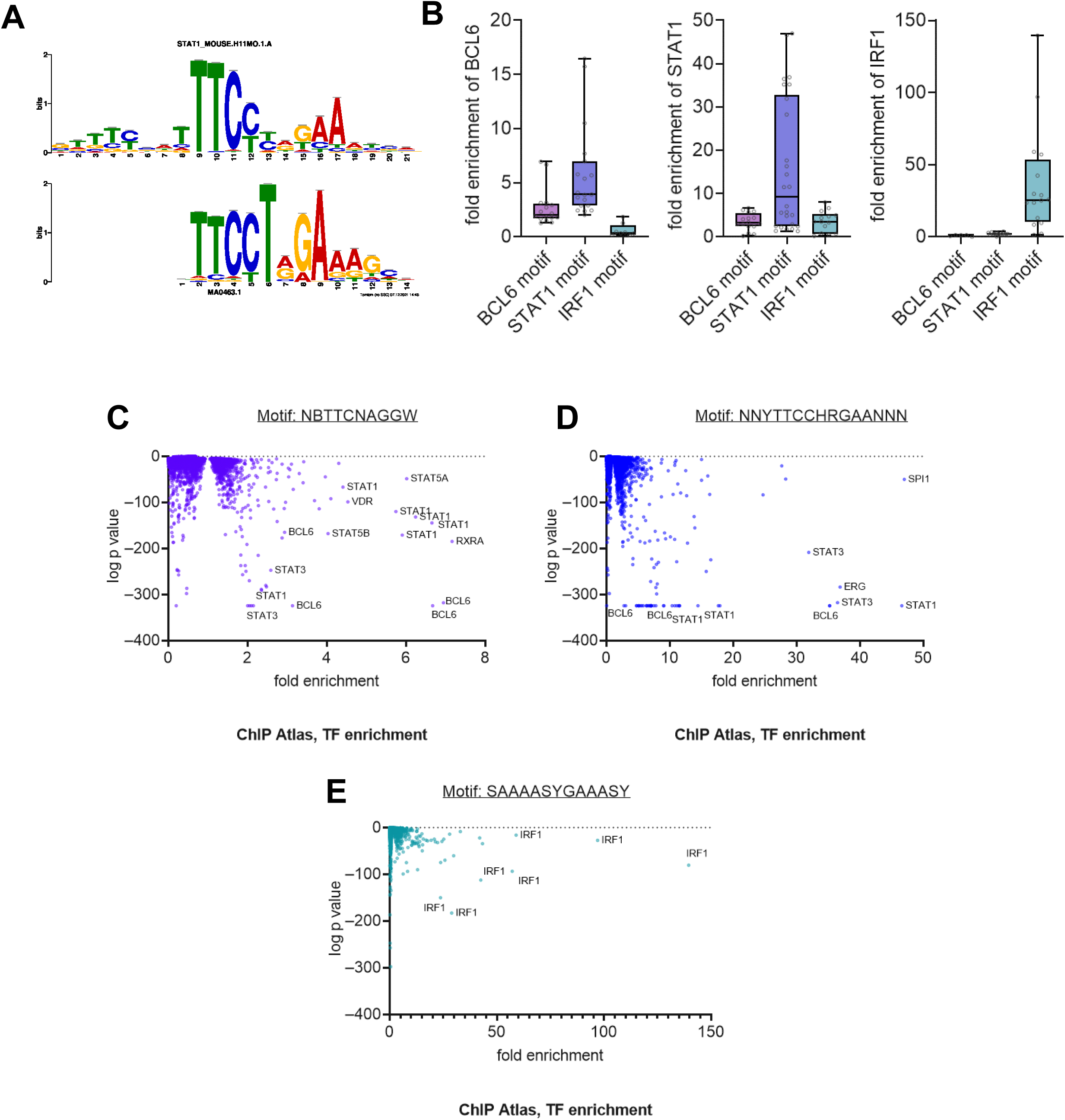
The DNA binding profiles of BCL6-STAT1 substantially overlap. A) The core STAT1 and BCL6 consensus motifs were aligned using JASPAR. B-E) Using the unbiased ChIP Atlas Enrichment Analysis, the degenerate consensus motifs for BCL6, STAT1, and IRF1 were queried against hundreds of public ChIP-Seq datasets for transcription factors with enriched binding to each DNA sequence. B) The distribution of each transcription factor’s fold enrichment at each consensus motif is shown in the box plots. C-E) The fold enrichment of all transcription factors in the ChIP Atlas database is plotted against the log p value, with annotations for the most highly enriched transcription factors binding to the (C) BCL6 motif, (D) STAT1 motif, and (E) IRF1 motif.

We performed an unbiased enrichment analysis using ChIP Atlas to identify transcription factors with enriched binding to the BCL6 consensus motif compared with a random permutation of the same dataset ^42^. Remarkably, other than BCL6 itself, the most highly enriched transcription factor binding to the NBTTCNAGGW motif (from MotifMap) was STAT1 (7.16-fold enrichment), in a ChIP dataset of K562 cells stimulated with IFNγ for 30min (GSM935487, from ENCODE GSE31477 ^41^). Indeed, of the top 50 TF Chip-Seq datasets with significantly enriched binding to this motif, BCL6 accounted for 8 (16%), STAT1 represented 13 (26%), and STAT3 was 10 (20%) (**Figure 4B, 4C**). We performed the same analysis for a second de novo BCL6 motif WRCTTTCBAGGRAT (from MotifMap), but only BCL6 itself and BCOR were significantly enriched for binding at this sequence (**Supplemental Figure 4B**). Reciprocally, we analyzed enrichment at the STAT1 motif NNYTTCCHRGAANNN, and the IRF1 motif SAAAASYGAAASY (**Figure 4B, 4D**). Among the top 100 most enriched TF binding to the STAT1 motif, BCL6 was identified 8 times (in 5 different cell types); STAT1 16 times; and other STATs 63 times. In contrast, BCL6 was not significantly enriched among TF binding to the IRF motif (**Figure 4B, 4E**). Overall, BCL6 fold enrichment at its own motif was 2.702±0.42-fold, while enrichment at the STAT1 motif was 5.70±0.97-fold, and at the IRF1 motif showed no enrichment 0.663±0.21-fold over control dataset. STAT1 showed significant enrichment at all three motifs: at the BCL6 motif 3.51±0.498-fold; at STAT1 motif, 15.91±3.04-fold; and at the IRF1 motif 3.56±0.64-fold (**Figure 4B**). Conversely, IRF1 had no enrichment for binding the BCL6 motif (0.918±0.23-fold) but did bind to the STAT1 motif (2.11±0.22-fold) as well as highly to its own motif (36.72±9.2-fold). These analyses demonstrate that BCL6 and STAT1 have significant overlap in genomic binding across different cell types and conditions, which is distinct from IRF recognition sequences.

Lastly, we mapped the STAT1 and BCL6 motifs along the human genome using the Sequence Motif Location Tool (molotool). The BCL6 consensus motif was entirely embedded within the IFN-responsive STAT/GAS motif. An example of overlapping motifs within the HLA-DPB1*01:01:01 gene is shown (**Supplemental Figure 4A**). In contrast, no similarity was found with IRF (IRE/ISRE motif).

Taking this evidence together, we hypothesized that BCL6 may function specifically as a specific regulator of STAT but not IRF transcription factors in the interferon response.

### BCL6 binding to interferon-response genes

The endothelial response to IFNγ is dominated by anti-viral, pro-adhesive, antigen presentation, and costimulatory gene expression. To further infer if BCL6 might regulate IFNγ response genes, we analyzed the BCL6 transcription factor motif distribution (MA0463.2, TFmotifView) across the genome, focusing on canonical interferon response genes, such as the MHC class II region (chr6:31437307-33447626 http://bardet.u-strasbg.fr/tfmotifview/?results=ezFBWpD8uTbPSB), PD ligands (chr9 5384082 5624107) and CXCL9-11 (chr4:75988158-76640626). The MHC class II region harbored numerous BCL6 DNA binding consensus motif sequences, both within genes, at the TSS and in intergenic regions. For example, one in intron 1 of *HLA-DRB1*, one in intron 1 of *HLA-DRB5*, another in intron 2 of *HLA-DPB1*, another midway between *DRB1* and *DQA1*, and another between *DQA1* and *DQB1*; but there were no motifs in or <2kb of DQ genes themselves (**Figure 4C**). Additionally, a BCL6 motif was found within *CIITA* intron 13, the last intron of *HLA-DOB*, and at multiple locations in and around the *IL15* gene.

Intriguingly, SNPs in STAT1 binding motifs, including in the HLA-A gene, are known to substantially affect gene expression after IFNγ stimulation ^47^. Because HLA is highly polymorphic, we asked whether the BCL6 motifs in the MHC class II region contained any common SNPs. The intragenic DPB1 motif and the intergenic DQA1/DQB1 motif are conserved. However, the intronic DRB1 motif has a common T/C SNP rs2874164 (eg. C in DRB1*01 and *04; T in DRB1*03); similarly the intronic DRB5 motif exhibits a SNP rs13894097 T/C variant (different position in the motif than for DRB1).

We next analyzed empirically defined BCL6 genomic binding peaks, using chromatin immunoprecipitation of BCL6 available from ENCODE in HepG2 and K562 cells ^41^. Using CistromeDB ^48^, we scanned top putative target genes encoding antigen presentation and costimulatory molecules that are previously defined as IFNγ-responsive in endothelium. We searched for BCL6 binding sites and identified significant peaks in the MHC class II region, particularly within the *CIITA* gene and between *HLA-DRB1* and *HLA-DQA1*, and within the HLA-DPA1 and –DPB1 genes (**Figure 4D**); at the TSS of the *TAP1* and *TAP2* genes; near the PD ligand genes *CD274* and *PDCD1L2*; and the cytokine/chemokine genes *IL15*, *CCL5* and *CX3CL1*. There was also a significant peak downstream of the *CXCL11-CXCL10-CXCL9* gene complex, although no peaks were found within those genes. In contrast, no BCL6 DNA binding peaks or motifs were found in or proximal to any HLA class I genes (**Figure 4D**); in the invariant chain gene *CD74*; *HLA-DMA*/*DMB*; *PSMB8, PSMB9*; *CD40*; or *TNFSF13B* (BAFF).

Intriguingly, within the MHC class II region, only a few highly significant peaks for BCL6 were found—about midway between the *HLA-DQA1* and -*DRB1* genes and at the *TAP1*/*TAP2* and *DMA*/*DMB* gene complexes. Next, we overlaid the public STAT1 genomic binding profile under IFNγ activation in HeLa cells ^49^. Consistent with the sequence similarity between BCL6 and STAT1 binding motifs, we found numerous highly significant overlapping BCL6-STAT1 sequences within target genes. For example, within *HLA-DPB1*, we identified 7 such directly overlapping sequences from intron 1 through intron 2. Some canonical interferon response genes, like within *MX1*, harbored a BCL6 recognition sequence, but did not exhibit BCL6 binding by ChIP-Seq in unstimulated HepG2 cells. This suggests that BCL6 localization may be cell type and/or context dependent, and reorganize across the genome under different cell states. Taken together, these analyses suggested that BCL6 might directly regulate STAT1-inducible antigen presentation and costimulatory molecules, and key cytokines involved in leukocyte activation.

### BCL6 overexpression suppresses IFNγ-induced HLA class II expression

Given this evidence, we next examined the functional role of BCL6 in endothelial cell activation. Endothelial cells were transduced with either control-mGFP or BCL6 ORF-mGFP to stably overexpress BCL6 (**Supplemental Figure 5A-5D**). Then, we tested the effects of BCL6 overexpression on IFNγ-induced antigen presentation molecules and chemokines in endothelium.

TeloHAEC stably overexpressing BCL6 or GFP were left unstimulated or activated with IFNγ for 24hr. Global gene expression was tested by RNA-Seq (n=2) (**Figure 6A**). Consistent with the flow cytometry data, BCL6 overexpression reduced HLA class II transcripts by 30-40% (**Figure 6C**). Additionally, chemokine mRNA *CXCL10* and *CX3CL1* were reduced. Conditioned supernatants from these experiments were assayed for secreted IP-10 (*CXCL10*) which confirmed that BCL6 overexpression reduced production of this chemokine.

**Figure 6.**
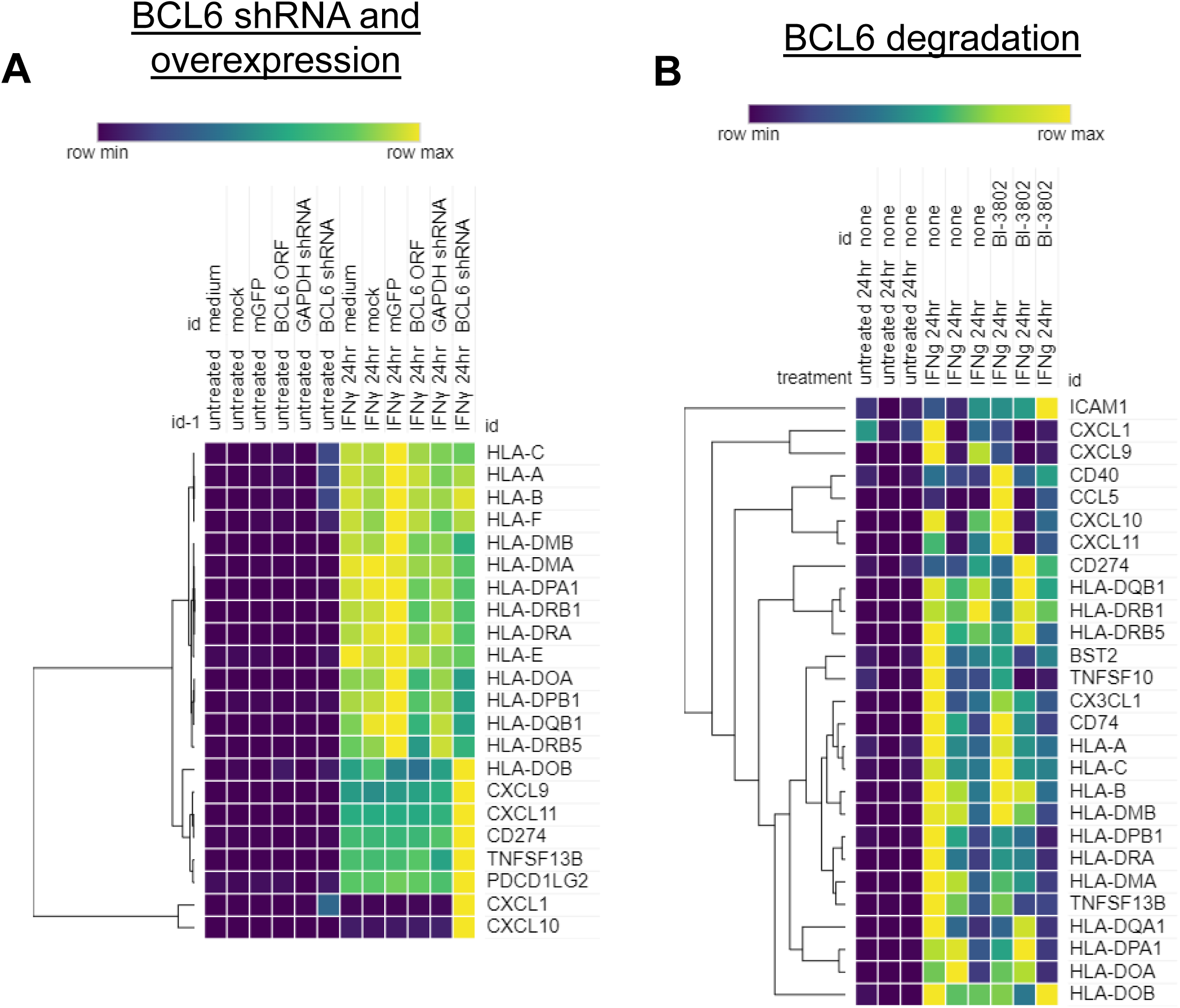
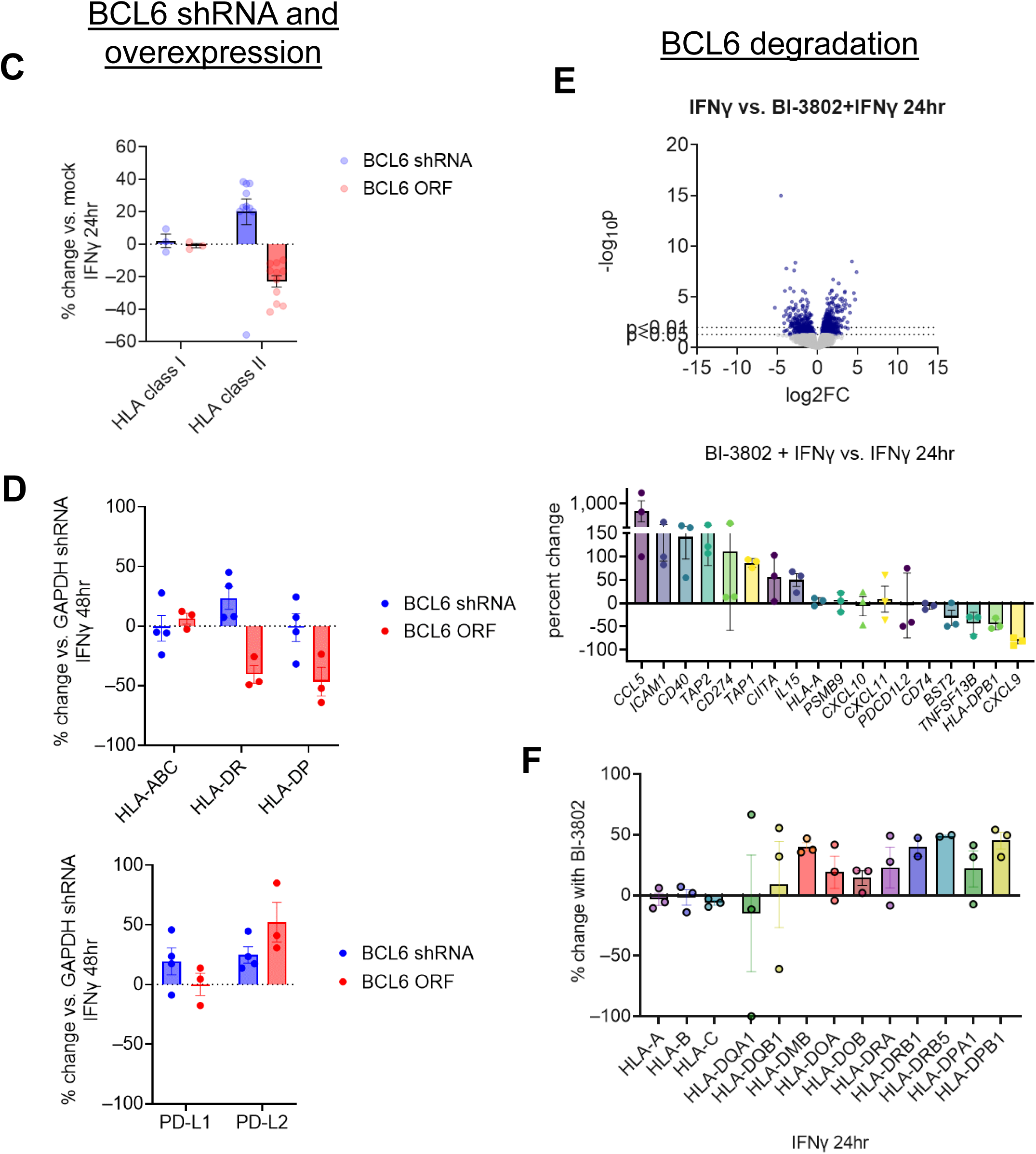
BCL6 knockdown and overexpression alters the endothelial response to IFNγ. A) TeloHAEC (n=2 technical replicates) stably transduced with the open reading frame of BCL6 or with GAPDH or BCL6 shRNA were stimulated with IFNγ for 24hr. Global gene expression was tested by RNA-Seq. Heat map shows the relative expression of major interferon inducible genes. B) Primary human aortic endothelial cells (n=3 biological replicates) were pre-treated with BI-3802 (100μM) to degrade BCL6 protein, followed by treatment with IFNγ for 24hr. Global gene expression was tested by RNA-Seq. Heat map shows the relative expression of major interferon inducible genes. C) The percent change in HLA gene expression under BCL6 knockdown and overexpression conditions is shown, comparing HLA class I genes (HLA-A, -B, and –C) to HLA class II genes (HLA-DRA, -DRB1, -DRB5, -DQA1, -DQB1, -DPA1, -DPB1, -DOA, -DOB, -DMA, -DOB). D) Primary human aortic endothelial cells were transduced with BCL6 shRNA or open reading frame, or pre-treated with BI-3802 to degrade BCL6 protein. Cell surface expression of HLA-ABC, -DR, and –DP was tested by flow cytometry after IFNγ treatment. E) Primary human aortic endothelial cells (n=3 biological replicates) were pre-treated with BI-3802 (100μM) to degrade BCL6 protein, followed by treatment with IFNγ for 24hr. Gene expression was measured by RNA-Seq. Volcano plot illustrates significantly differentially expressed genes comparing IFNγ alone to BI-3802 and IFNγ combined. Waterfall plot shows the percent change in top differentially expressed genes comparing IFNγ alone to BI-3802 and IFNγ combined. F) The percent change comparing IFNγ alone to BI-3802 and IFNγ combined in HLA class I and HLA class II genes is graphed.

Primary HAEC transduced with BCL6 (12.5-100 MOI) or GFP-alone (25 MOI) lentivirus were treated with IFNγ for 24-48hr. Cell surface expression of PD ligands and HLA molecules was measured by flow cytometry. Overexpression of BCL6 reduced HLA-DR (−49.15±6.1%) and HLA-DP (−46.56±9.8%) protein expression by nearly half compared with IFNγ under control transduction conditions (**Figure 6D**), but did not alter HLA class I (+5.71±9.02%) expression after IFNγ treatment. Suppression of HLA-DR and HLA-DP was dose responsive, where 100 MOI yielded the strongest inhibitory effect, and 25 and 12.5 MOI showed very little inhibition. These results demonstrate that BCL6 functions to selectively repress IFNγ-inducible expression of HLA-DR and HLA-DP, but not HLA class I.

### BCL6 loss enhances an interferon signature in endothelial cells

We then performed reciprocal experiments testing depletion of BCL6 using two approaches. Treatment of cells with the small molecule BI-3802 causes rapid BCL6 degradation ^50, 51^, and gene expression changes in B cell lines. As a parallel approach, we transduced endothelium with GAPDH or BCL6 shRNA using lentiviral vectors. To test whether loss of BCL6 affected the endothelial type II interferon response, we examined global gene expression in immortalized TeloHAEC transduced with shRNA against BCL6 (**Figure 6A, 6C, 6D**), and primary human aortic endothelial cells pre-treated with BI-3802 (**Figure 6B, 6E, 6F**), to deplete BCL6 prior to IFNγ stimulation.

In untreated EC, many of the genes that were most highly upregulated by BCL6 shRNA or BI-3802 depletion were canonical interferon response genes (**Figure 6A, 6B, 6E**). In particular, expression of *STAT1, MX1, MX2, OAS1, OAS2, ISG15,* and the effector adhesion genes *CXCL11, CXCL10, BST2* was increased in untreated BCL6 shRNA TeloHAEC. Additionally, BCL6 depletion, but not GAPDH shRNA, increased *IFI35*, *IFI44*, *IFI44L*, *IFI6*, *IFIH1*, *IFIT1*, *IFIT1B*, *IFIT2*. Of 595 genes were increased by IFNγ treatment at 24hr (CPM >2.0, >2-fold) 132 genes were increased >50% by BCL6 shRNA, 8 of which were very highly (>5-fold) increased: *CXCL10*, *CXCL2*, *CXCL3*, *BIRC3*, *MX2*, *NUAK2*, *RSAD2* and *UBD*. In particular, after IFNγ stimulation, BCL6 shRNA further increased transcript abundance of the CXCR3 chemokines *CXCL9* (2.1-fold), *CXCL10* (31.9-fold), and *CXCL11* (21-fold).

Using flow cytometry, we confirmed that shRNA against BCL6 slightly but consistently augmented IFNγ-induced (48hr) HLA-DR (+16.36±7.02%) and PD-L2 (+18.15±2.3%) cell surface expression compared with control GAPDH shRNA transduced conditions (n=3) (**Figure 6D**). Additionally, BCL6 shRNA augmented expression of PD-L2, but not PD-L1 (**Figure 6D**).

We then verified the effect of BCL6 loss in primary HAEC treated with the degrading inhibitor BI-3802. 450 and 301 genes were significantly increased and decreased (±2-fold, p<0.05), respectively, by BI-3802+IFNγ compared with IFNγ alone (**Figure 6E**). As shown in the waterfall plot, top genes that were significantly increased by BI-3802 were *CCL5, ICAM1, CD40, TAP2, CD274, CIITA,* and *IL15*. On the other hand, BI-3802 degradation of BCL6 caused significant downregulation of *BST2, TNFSF13B, HLA-DPB1* and *CXCL9*. BI-3802 also augmented IP-10 secretion by primary HAEC (4.95±5.4-fold, n=3), similar to the effect of BCL6 shRNA (**Figure 6E, 6F**). Depletion of BCL6 with either shRNA or BI-3802 also substantially increased CCL5 (RANTES) production by IFNγ-treated endothelium.

These results show that loss of BCL6 substantially alters the IFNγ-induced transcriptome of cardiac endothelium, primarily increasing expression of ISGs and HLA class II molecules.

### Effect of pharmacological targeting of the BCL6-BTB domain on response to IFNγ

A major mechanism of action by which BCL6 regulates transcription of target genes is through association with corepressors (SMRT/NCOR, BCOR) that recruit histone deacetylases (HDAC3; HDAC4, 5, 7) and other epigenetic modifiers ^21, 23, 24, 52^. In untreated endothelial cells, BCOR was moderately expressed, while NCOR1 and SMRT/NCOR2 were highly expressed; but none were changed by IFNγ treatment.

Several small molecule inhibitors have been developed that target the BTB domain of BCL6 to block association with corepressors BCOR and NCOR/SMRT. These include BI-3812, 79-6, FX1, and the peptidomimetic RI-BPI ^38, 50, 53, 54^. FX1 is a high potency derivative of 79-6. To understand the BTB domain-dependence of BCL6 action in the IFNγ pathway in endothelium, we pre-treated endothelial cells with 79-6, FX1, and BI-3812 and tested IFNγ induced gene expression.

Surprisingly, targeting the BTB domain of BCL6 both enhanced and suppressed IFNγ-induced gene expression in endothelium (**Figure 7A, 7B**). Among IFNγ-induced genes, FX1 downregulated *HLA-DOA, HLA-DRA, CXCL10, CXCL9, CD74, CX3CL1, CXCL11, SEPTIN4* and *HLA-DRB1*. Similarly, top genes decreased by BI-3812 were *CCL5, ICAM1, TAP2, CD40, CD274, IL15*, *CD74, BST2, CXCL10, CXCL11, TNFSF13B, HLA-DPB1* and *CXCL9*. (**Figure 7C, 7D**). Both FX1 and BI-3812 significantly inhibited expression of *CXCL9*, *CXCL10*, *VCAM1*, *TNFSF13B*, *PDCD1LG2*, *CD74*, *HLA-DPB1*, *HLA-DRA*, *MX1*, *MX2* and *ISG15*. A subset of genes were inhibited by the presence of FX1 but were not significantly changed by BI-3812: *CIITA*, *CX3CL1*, *HLA-DQB1*, *PSMB8*, *OAS2*, *IDO1*, *HLA-F* (↓). On the other hands, some genes were augmented by both inhibitors, including *TAP1, TAP2, HLA-F, CD274, BCL6, MICA, HLA-E, CSF1, IRF2,* and *NKX3-1*. As with overexpression, here was no significant change in IFNγ-induced expression of *CD40, HLA-A, HLA-B, HLA-C, IL15, ICAM1,* or *BST2*. Importantly, similar patterns of gene expression were observed with the two BTB inhibitors, although FX1 was notably more potent, confirming the mechanism of action.

**Figure 7.**
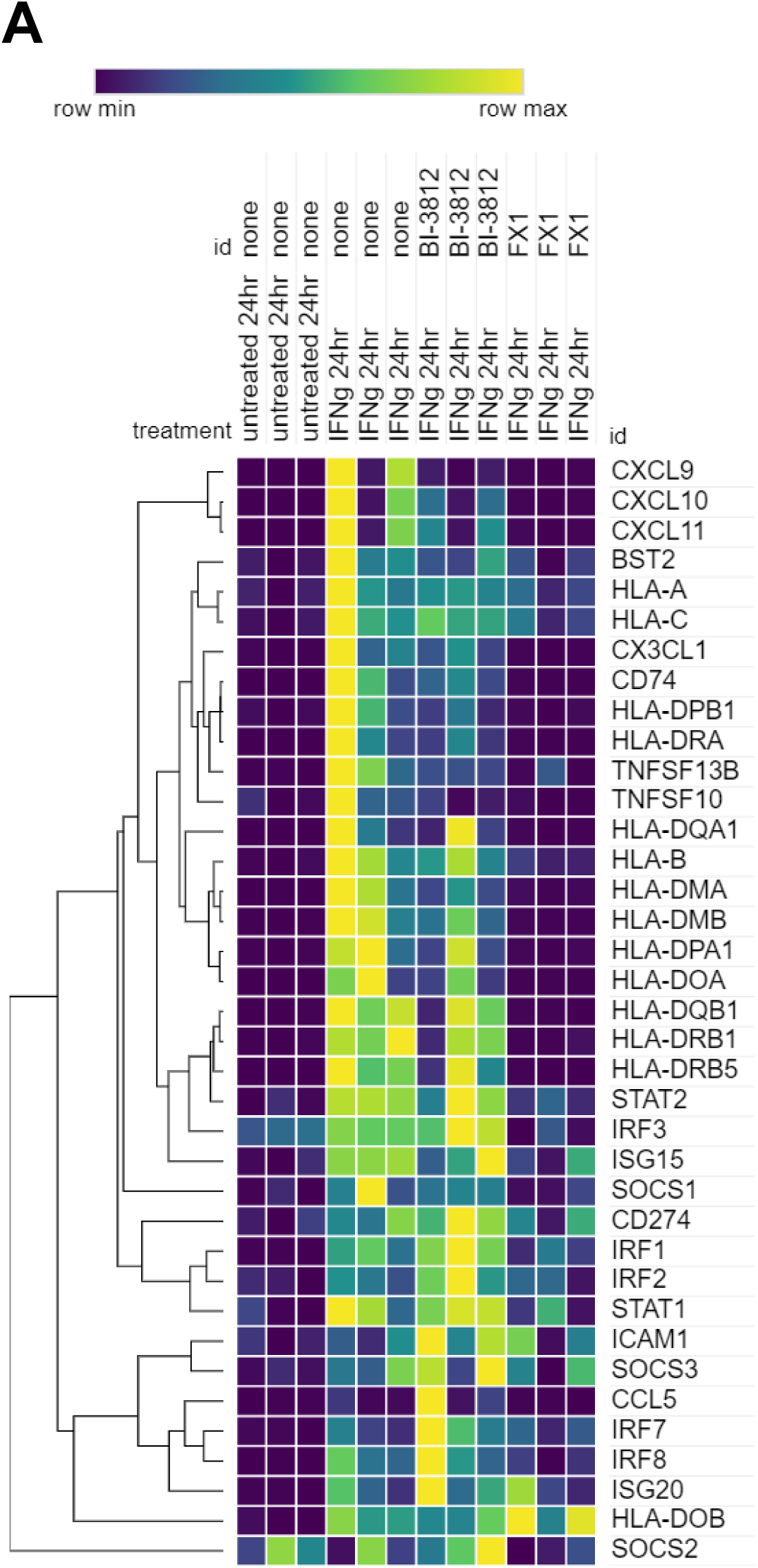
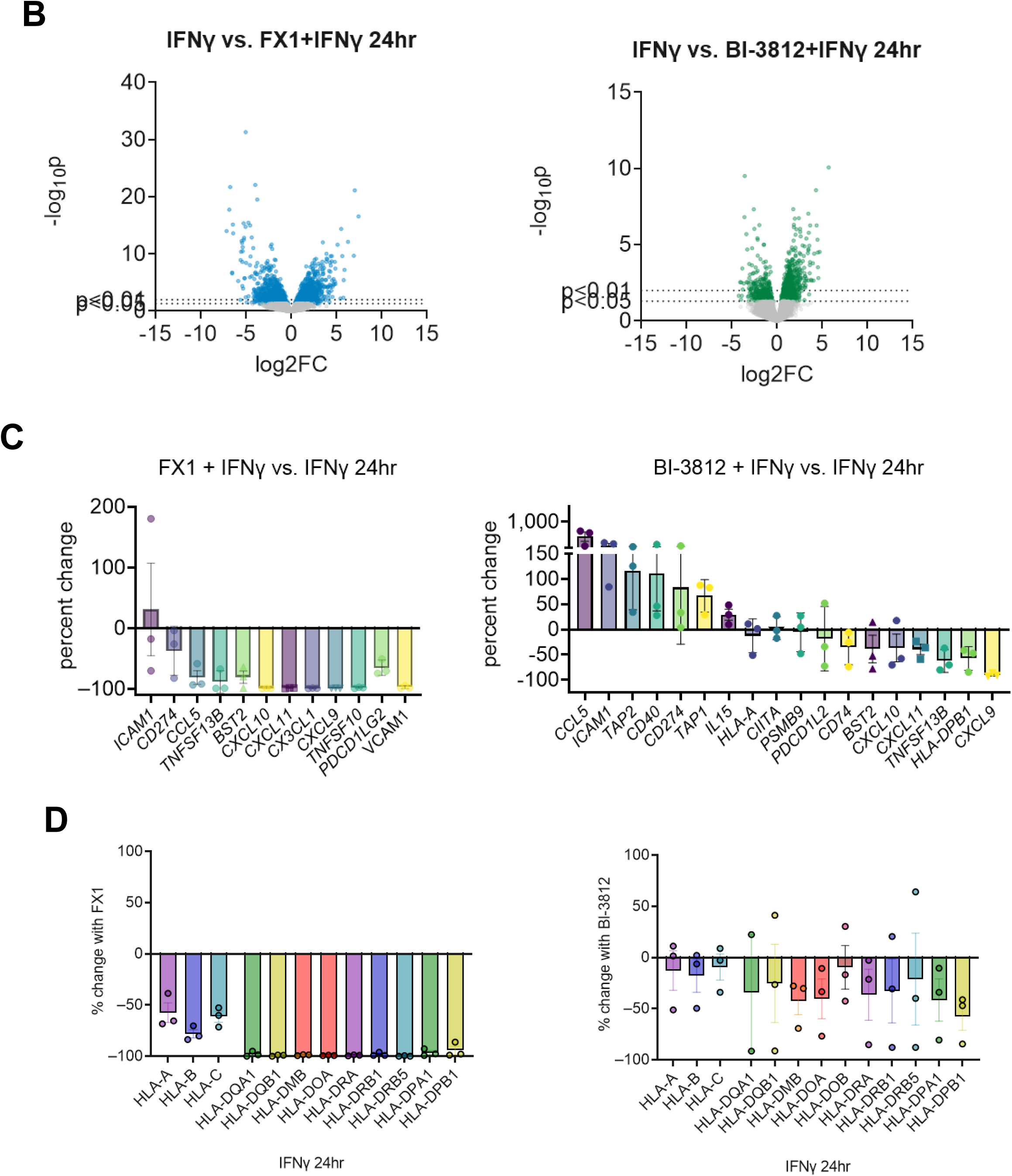
BCL6-BTB antagonists suppress much of the IFNγ response in endothelial cells. A) Primary human aortic endothelium (n=3 biological replicates) were pre-treated with FX1 or BI-3812 (100μM), followed by IFNγ for 24hr. Gene expression was tested by RNA-Seq. Heat map shows relative expression of major interferon response genes. B) Volcano plots illustrate significantly differentially expressed genes comparing IFNγ alone to BI-3812 or FX1 and IFNγ combined. C) Waterfall plot shows the percent change in selected gene expression comparing IFNγ alone to BI-3812 and IFNγ combined. D) The percent change comparing IFNγ alone to BI-3812 or FX1 and IFNγ combined in HLA class I and HLA class II genes is graphed.

IPA analysis was performed to identify differentially expressed gene modules. Comparing IFNγ alone to IFNγ+FX1, top canonical pathways included antigen presentation (p=1.17e-13), interferon signaling (p=1.16e-09) and PD1/PD-L1 cancer immunotherapy (p=2.86e-08); top regulator effect networks comprised “activation of leukocytes”.

We verified the effects of BCL6-BTB domain inhibitors on expression of HLA and costimulatory molecules, and secretion of chemokines and cytokines. Pre-treatment of endothelial cells with FX1 or BI-3812 did not affect mRNA or protein expression of CD40, PD-L1 (*CD274*), *FAS* or *IL15* (**Figure 8A**). However, IFNγ-induced expression of BAFF (*TNFSF13B*) and PD-L2 (*PDCD1LG2*) were significantly suppressed by BTB domain antagonists (**Figure 8A, 8B**). Additionally, while HLA class I expression was unchanged after IFNγ for 24hr or 48hr, in contrast, BTB inhibitors significantly inhibited mRNA and cell surface expression of HLA class II molecules HLA-DR (24hr), and HLA-DP (24hr) (**Figure 9A, 9B**). The effect of FX1 and BI-3812 on HLA class II expression was dose responsive, but even the highest dose tested had no effect on HLA class I expression (**Supplemental Figure 9**). Interestingly, other HLA class I related antigen presentation genes TAP1, TAPBP and TAP2 were also unchanged by BCL6-BTB antagonists. As a control, the low potency derivative of FX1, 79-6, had little to no effect on HLA or PD ligand expression (**Figure 8B, 9B, Table 2**).

**Figure 8.**
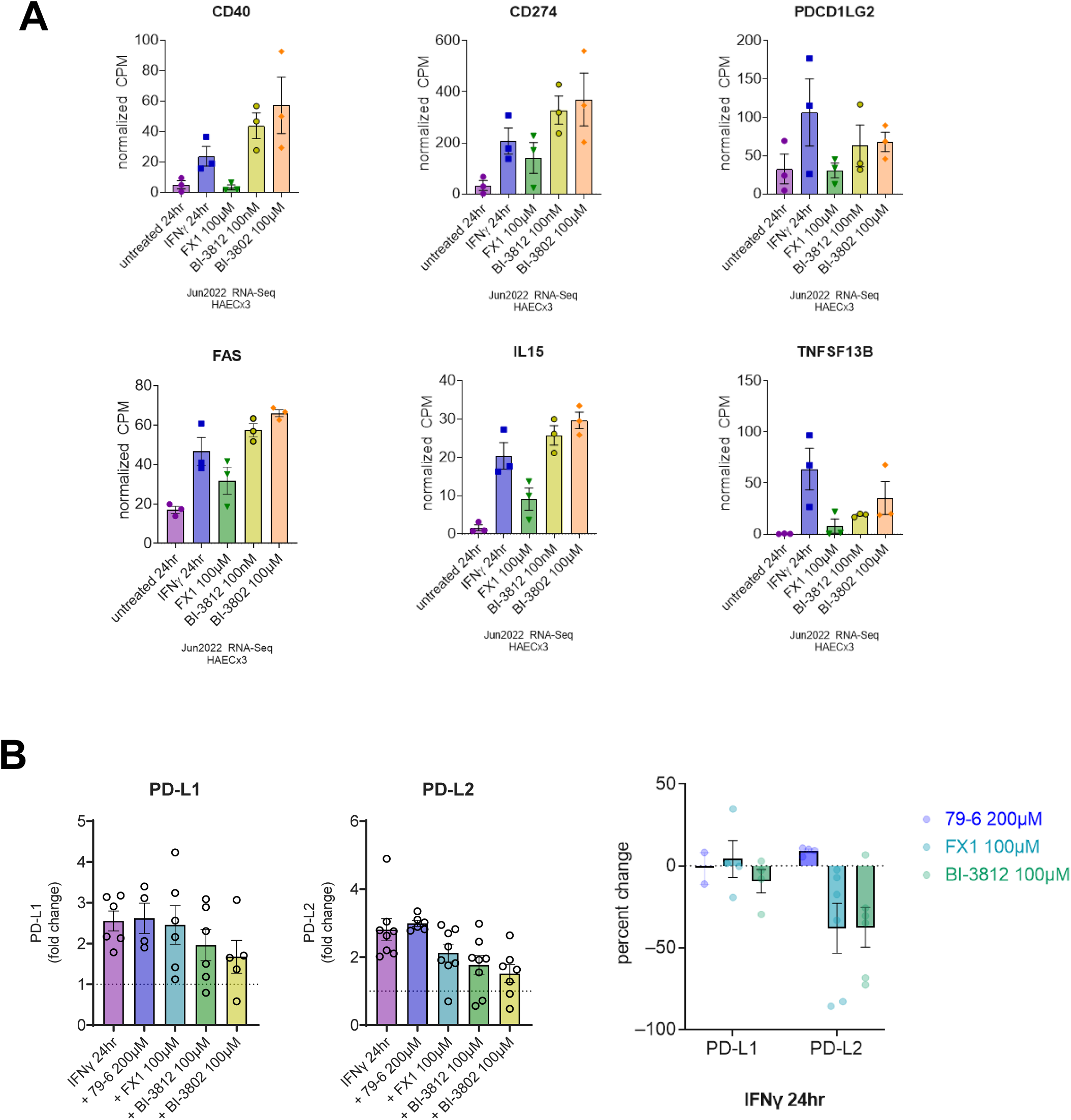
Effect of BCL6 on IFNγ-induced expression of costimulatory molecules. A) Primary human aortic endothelial cells (n=3 biological replicates) were pre-treated with FX1 or BI-3812 (100μM) followed by IFNγ for 24hr. Gene expression was measured by RNA-Seq. Graphs show normalized CPM for costimulatory gene expression. B) Primary human aortic endothelial cells (n≥5 biological replicates) were pre-treated with 79-6 (200 μM), FX1 or BI-3812 (100μM) followed by IFNγ for 24hr. Cell surface expression of PD ligand proteins was measured by flow cytometry. Fold change compared to untreated parallel control cells is graphed on the left. Percent change is graphed on the right.

**Figure 9.**
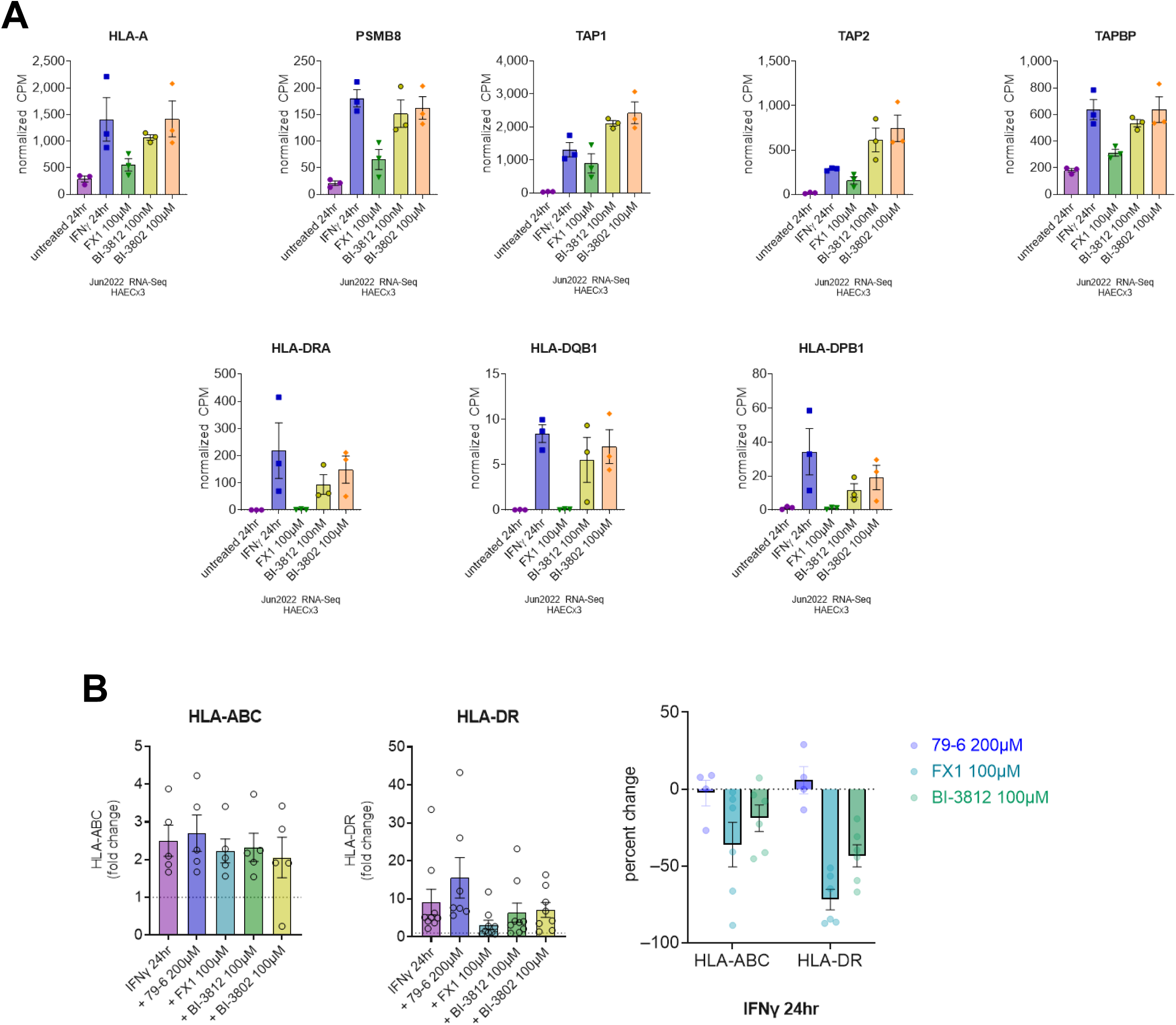
Effect of BCL6 inhibitors on IFNγ-induced HLA expression in endothelial cells. A) Primary human aortic endothelial cells (n=3 biological replicates) were pre-treated with FX1 or BI-3812 (100μM) followed by IFNγ for 24hr. Gene expression was measured by RNA-Seq. Graphs show normalized CPM for HLA class I-related and HLA class II gene expression. B) Primary human aortic endothelial cells (n≥5 biological replicates) were pre-treated with 79-6 (200 μM), FX1 or BI-3812 (100μM) followed by IFNγ for 24hr. Cell surface expression of HLA proteins was measured by flow cytometry. Fold change compared to untreated parallel control cells is graphed on the left. Percent change is graphed on the right.

**Table 2.** Percent inhibition of protein expression (IFNγ 24hr).

|  | **HLA I** | **HLA-DR** | **PD-L1** | **PD-L2** |
| --- | --- | --- | --- | --- |
| **79-6** | 8.17±2.58 | 72.18±46.0<br>4 | 10.48±8.89 | 4.45±15.11 |
| **FX1** | -38.84±17.55 | -<br>78.82±9.83 | 25.74±45.0<br>1 | -<br>31.97±22.00 |
| **BI-3812** | -19.20±10.87 | -<br>53.82±12.4<br>6 | -<br>8.73±28.92 | -<br>60.99±18.65 |
| Table 2. Percent inhibition of protein expression (IFNγ 24hr). | | | | |

Among pro-adhesive genes, FX1 and BI-3812 (100μM) both significantly reduced the production of MIG (*CXCL9*), IP-10 (*CXCL10*), and I-TAC (*CXCL11*) by endothelial cells treated with IFNγ for 24hr (**Figure 10A, 10B**). On the other hand, the BCL6-BTB inhibitor BI-3812 augmented of RANTES (*CCL5*), while control 79-6 had no effect on any chemokines tested (**Figure 10B**). Additionally, IFNγ-induced *VCAM1* expression by endothelial cells was completely blocked by both BTB inhibitors, while *ICAM1* and *BST2* were relatively unaffected (**Figure 10A**).

**Figure 10.**
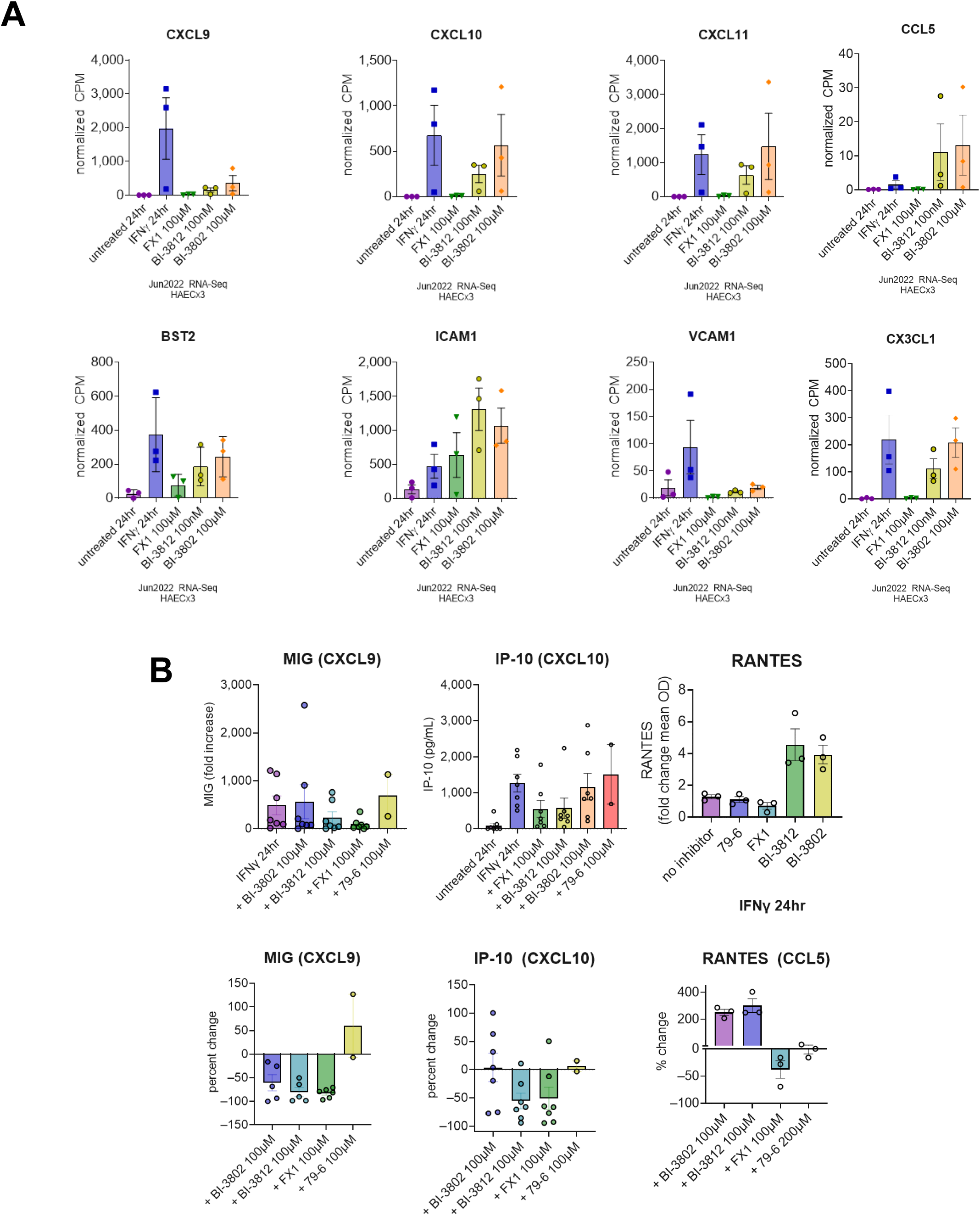
Effect of BCL6 on IFNγ-induced expression of pro-adhesion molecules and chemokines. A) Primary human aortic endothelial cells (n=3 biological replicates) were pre-treated with FX1 or BI-3812 (100μM) followed by IFNγ for 24hr. Gene expression was measured by RNA-Seq. Graphs show normalized CPM for chemokine and adhesion molecule gene expression. B) Conditioned medium from primary human aortic endothelial cells (n≥5 biological replicates) pre-treated with 79-6 (200 μM), FX1 or BI-3812 (100μM) followed by IFNγ for 24hr were tested for secreted chemokines by ELISA.

These results show that antagonism of the BTB domain of BCL6 potently and selectively inhibits IFNγ response gene expression, an effect which was similar to, but stronger than, BCL6 overexpression.

### BCL6 inhibitors effect on STAT activation

To understand the mechanism by which BCL6 affects the IFNγ response in endothelium, we asked whether BCL6 inhibitors directly affected activity of the major IFNγ-activated transcription factor STAT1. First, we probed STAT1 localization in endothelium either pre-treated with BCL6 inhibitors or transduced with shRNA or open reading frame of BCL6 prior to IFNγ stimulation. STAT1 staining was diffuse and faint in untreated cells, but became strongly nuclear after IFNγ 30min. BCL6 overexpression and knockdown had no effect on STAT1 localization after IFNγ stimulation (**Figure 11A**).

**Figure 11.**
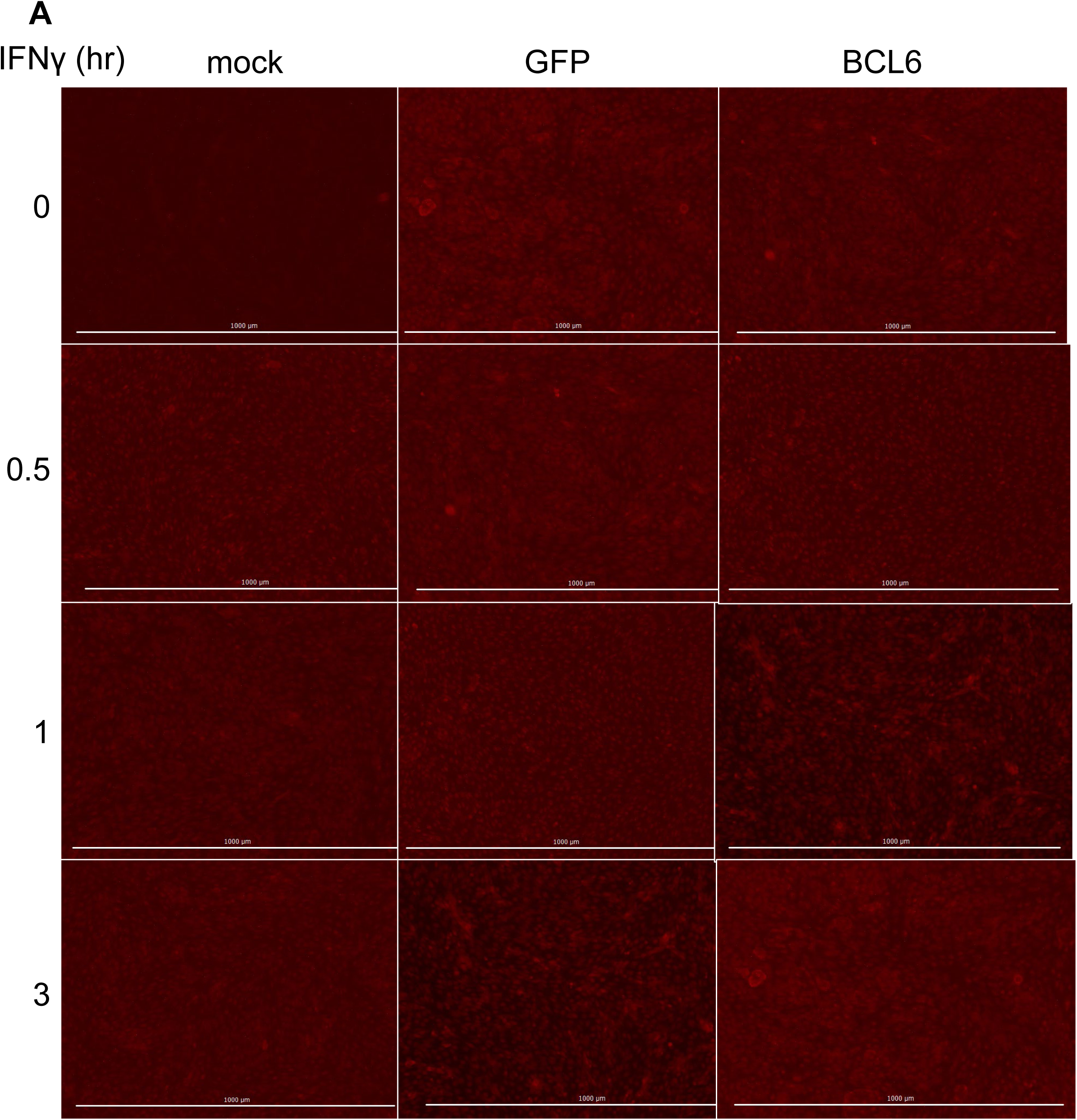
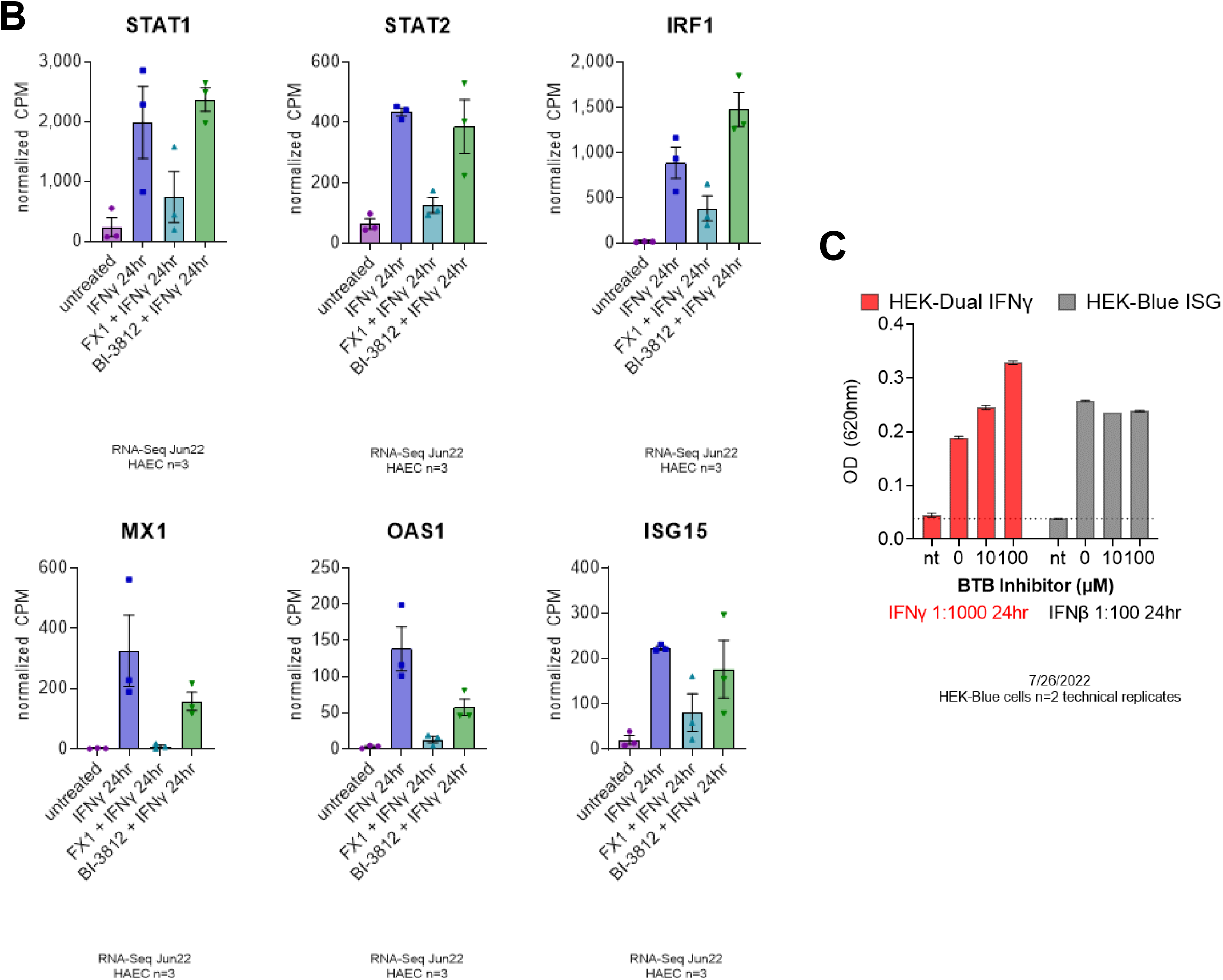
Effect of BCL6 inhibitors on STAT1 dependent signaling. A) TeloHAEC transduced with no lentivirus, GFP alone, or BCL6 overexpression were treated with IFNγ for 0.5-3hr. Immunofluorescence micrographs show STAT1 intracellular localization in IFNγ-treated TeloHAEC over time. B) Normalized gene expression from RNA-Seq data showing STAT, IRF, and ISG transcript counts in primary human aortic endothelial cells (n=3). C) BTB antagonism of BCL6 affects STAT/GAS-dependent transcription but not IRF/ISRE. Reporter HEK-Blue cells were pre-treated with increasing concentrations of FX1 or BI-3812, and treated with IFNγ or IFNβ for 24hr. Reporter activity was measured by SEAP.

Further, BCL6 overexpression and knockdown had no effect on increased expression of IRF1 or STAT1 transcription factors themselves, nor other IRFs or STATs except STAT2 which was increased with BCL6 shRNA + IFNγ (**Figure 11B**). In addition to the inflammatory genes described above, we were interested in the effect of BCL6 on canonical interferon signaling genes, many of which are involved in the anti-viral response. *TRIM14, OAS1/2/3, OASL, ISG15, ISG20, MX1* and *MX2* were all increased basally in BCL6 shRNA transduced cells, and very highly upregulated after IFNγ compared with GAPDH shRNA. Conversely, FX1 and BI-3812 both suppressed IFNγ-induced expression of *OAS1/2/3, ISG15, IDO1* (**Figure 11B**). However, there was no change in other interferon-inducible anti-viral genes *ADAR*, *TRIM21*, or *TRIM22*.

Then, we tested the reporter cell lines HEK-Dual IFNγ and HEK-Blue ISG, HEK293 cells stably transfected with SEAP reporter driven by either a GAS motif exclusively responsive to STAT1, or as a control, an ISRE/ISG54 motif only responsive to type I IFNs via IRFs (**Supplemental Figure 10**). The BTB domain inhibitors FX1 and BI-3812 dose-dependently increased STAT1 transcriptional activity (**Figure 11C**). On the other hand, FX1 and BI-3812 had no effect on type I IFNβ-induced IRF transcriptional activity, suggesting a specificity for STAT1-mediated transcription (**Figure 11C**).

Together, these results demonstrate that BCL6 directly affects STAT1 transcriptional activity.

## Discussion

Although transplantation is a life-saving therapy for tens of thousands of patients with end-stage organ failure each year, immune-mediated rejection is a significant cause of morbidity and graft failure. Organ transplant rejection is a serious complication arising from recipient alloimmune activation, causing donor vascular and parenchymal inflammation ^55^. Current immunosuppression regimens impair adaptive alloimmunity to prevent rejection, but adversely impact protective immunity. Even still, acute rejection occurs in up to a third of heart transplant recipients in the first year post-transplant ^56^. Prior acute rejection episodes are a significant risk factor for later development of allograft vasculopathy and graft loss ^57–59^. As a result, the half-life of heart transplants is much shorter than recipient life expectancy, ranging from 5 to 15 years. During rejection, recipient alloreactive leukocytes and antibodies directly target the donor endothelial cells lining the graft blood vessels. The graft is an active participant in the rejection process, but current therapies do not target the donor organ. Rejecting allografts exhibit both interferon (IFN) and endothelial gene expression signatures, and IFNγ activation of endothelium promotes inflammation by upregulating molecules that enhance leukocyte recruitment and shape alloreactive T cell activation. However, the field lacks knowledge of the intrinsic regulators of this process, and consequently efficacious therapies to locally dampen graft rejection are lacking.

Our results show that the transcriptional repressor BCL6 is increased in rejecting human heart transplants, is IFNγ inducible, and in turn suppresses a portion of the IFNγ gene programming in endothelial cells. To our knowledge, this is also the first report of BCL6-mediated regulation of MHC class II expression in any cell type. Further, while BCL6 regulation of PD ligands and CXCL chemokines has been reported in B cells and macrophages ^33, 60^, our study is the first to demonstrate this effect in endothelial cells. Specifically, we hypothesize that BCL6 functions as a transcriptional repressor of IFNγ/STAT1 to dampen pathogenic endothelial gene expression that occurs during transplant rejection.

### Transplant Rejection

Transplant rejection is a consequence of alloimmunity, in which the recipient adaptive immune system recognizes genetic variants in the donor as foreign. The most prominent target of alloimmunity is the major histocompatibility complex (HLA system in humans), which is highly polymorphic. Among the 12 loci in the HLA complex, HLA-DQ mismatches are considered to be the most detrimental to allograft survival across organs and are the most frequent targets of anti-donor alloantibodies ^61^. Despite this overwhelming clinical knowledge that HLA-DQ is most important to transplant outcomes, the underlying biology is not at all defined. Our data surprisingly show that HLA-DR and HLA-DP are upregulated by IFNγ-treated endothelial cells more quickly compared with HLA-DQ, at both the transcript and protein levels. These intriguing results suggest that the mechanisms of HLA-DQ expression and regulation may be distinct from those of other HLA class II genes.

In addition to the immunogenic HLA molecules, endothelial cells conditionally express a wide array of costimulatory molecules and cytokines. Endothelial PD-L1+HLA-DR+ expression by flow cytometry from human heart transplants, inversely correlated with CD8+ T cell infiltration during rejection ^12^. In mice, PD-L1 but not PD-L2 was critical for inhibitory signals that prevent rejection. Donor endothelial deletion of PD-L1 caused accelerated rejection and graft loss in mouse heart transplant model ^12^.

### BCL6 Transcriptional Repressor

BCL6 is a transcriptional repressor critical for germinal center formation, Tfh cell lineage specification, macrophage activation, and regulation of cytokine responses ^20, 21, 23, 24, 28, 62–64^. BCL6 contains zinc finger (ZF) DNA binding domain; a linker/RD2 region; and a BTB domain through which it complexes with corepressors ^24, 65^. BCL6 interacts with histone deacetylases through the RD2 and ZF domains ^52, 66, 67^, and indirectly via BTB domain-associated corepressors ^68^.

BCL6 is IFNγ/STAT-inducible in several cell types ^26–31^. Its loss derepressed interferon response and JAK/STAT gene modules, in part through competition with STATs for GAS DNA sequences ^27, 69–73^. IFNγ also causes chromatin remodeling to reshape the enhancer landscape of inflammatory genes ^74^. Work from the Evans lab showed that JAK-STAT KEGG pathways were significantly affected in macrophages by RI-BPI ^23, 24^, supporting a substantial role for BCL6 in interferon responses. Additionally, BCL6 suppressed IFNγ-induced genes such as PD-L1 and PD-L2 in germinal center B cells, and *CXCL9* in the liver ^33, 75^. In adipocytes, nearly as many genes were downregulated in *Bcl6* knockouts as upregulated ^25^, pointing to a dual effect of BCL6 on gene expression. The Crotty group has proposed that BCL6 also carries out “repression of repressors” such as miRNA or alternative lineage transcription factors, raising an alternative mechanism by which BCL6 selectively steers cytokine responses ^20^.

BCL6 partly exerts its repressive function through chromatin regulation via its BTB domain ^76^. Each of BCL6’s domains has independently been shown to independently confer repressive function ^21, 65–67, 76, 77^. However, the comparative contributions of each functional domains and requirements for corepressors in endothelium remains to be determined.

The BTB domain dependence of BCL6 may be cell type dependent, since B cells but not T or macrophage phenotypes were altered by a BTB mutation ^66, 76^, and mice with BTB mutant domain of *Bcl6* do not exhibit the lethal inflammatory syndrome of *Bcl6* knockout mice ^66^. Also, treatment of mice with BTB antagonists diminishes rather than exacerbates inflammation and organ damage in mouse models ^78, 79^. In hepatocytes and macrophages, roughly 20% of the BCL6 genomic binding sites were unique to BCL6 and did not overlap with SMRT or NCOR^23, 25^. In B cells, H3K27Ac at BCL6-bound enhancers lacking SMRT was not affected by a BTB domain inhibitor ^80^. The ZF region of BCL6 alone was sufficient to exert transcriptional repressor activity at least in a highly manipulated *in vitro* system ^52^.

In addition to complexing through the BTB domain, BCL6 can physically associate with class II HDACs (4, 5, 7) through its ZF domain (so independent of BTB/corepressors). Further, mutation or antagonism of the BTB domain does not reduce binding of BCL6 to its DNA targets ^38, 79^, and several groups have surmised that BTB inhibitors enhance BCL6 DNA binding ^38, 79^, to explain their anti-inflammatory/increased repressive effect. Substantial data exist in the literature describing several discrete mechanisms of action of BCL6 in other cell types. These include direct regulation and competition or cooperation with other transcription factors ^15, 21, 27^, indirect regulation by controlling gene accessibility through chromatin remodeling ^52, 64, 67^, and interactions with other repressors such as microRNAs (miRs) to secondarily affect gene expression post-transcription ^81, 82^. In B cells, BCL6 preferentially localized to less accessible chromatin regions and negatively correlated with transcriptional activity of nearby genes ^83^. BCL6 forms homodimers through the BTB domain. At the interface of the dimers, corepressors such as NCOR/SMRT bind and subsequently engage HDACs. This interaction decreases H3K27Ac, and halts transcription in the hematopoietic lineage ^25, 65, 68, 80, 84^. Nevertheless, as elegantly shown in GC B cells, Tfh cells and macrophages in the mouse, the BTB domain is surprisingly dispensable for repression of target genes depending on the context. For example, BCL6 instead directly antagonizes STAT transcription factor binding ^76^.

Based on this literature, we posit that BTB domain antagonists (BTBi) that dissociate corepressors actually enhance BCL6 dimerization and DNA binding, as shown in ^38, 79^, thereby augmenting its suppressive effects. We propose that this stabilizes its competition with STAT1 to cause more potent gene repression.

### BCL6 Pharmacological Targeting

Standard immunosuppression (CSA, TAC) did not affect IFNγ-induced HLA-DR expression on endothelium ^85^. However, our results demonstrate that overexpression or pharmacological targeting of BCL6 effectively suppressed HLA-DR and HLA-DP expression, with had variable effect on HLA-DQ.

The BCL6 BTB domain has low homology with other proteins, enabling highly specific targeting of BCL6’s corepressor interface ^38, 50^. We had expected to find that BTB domain inhibitors would derepress gene programming but were surprised to find that inhibition of the BTB domain caused profound suppression of many IFN response genes. Although causing efficient dissociation of NCOR/BCOR, BTB domain inhibitors do not dissociate BCL6 from target DNA, and apparently increase binding by 10- to 40-fold ^38^. Another study showed in cancer stem cells also found that FX1 treatment increased BCL6 binding at target genes even while abrogating association with BCOR ^86^. And, in a mouse models of LPS-induced sepsis, GVHD, and autoimmunity, treatment of mice with FX1 unexpectedly protected from severe tissue inflammation ^79, 87^.

In adipocytes, BCL6 basally occupied target genes and was displaced by STAT5 and the chromatin modifier p300 upon growth hormone stimulation ^88^. The Melnick lab found that the DNA binding domain, but not the BTB domain, of BCL6 was required for suppression of cytokine response genes by macrophages, and they similarly concluded that direct competition between BCL6 and STATs was responsible for BCL6 repressive activity, at least in innate immune cells ^76^. This model suggests that BCL6 directly represses STAT target genes under basal conditions, and is released to permit STAT-driven transcription in stimulated cells. Taken together, we hypothesize that BTB antagonists stabilize BCL6 binding to repressed target genes, which blocks STAT1-driven gene expression under IFNγ stimulation. Experiments are ongoing to define the overlap between the BCL6 and STAT1 cistromes in endothelial cells, and determine how BCL6 occupancy changes under IFNγ stimulation, affects chromatin accessibility, and alters STAT1 binding to target genes.

In summary, our results demonstrate that targeting BCL6 has a potent anti-inflammatory effect on IFNγ-induced gene expression in endothelial cells. Taking the literature and our results together, we propose a mechanistic model (**Figure 12**), in which BCL6 directly binds to STAT1 DNA recognition sequences to repress target genes. Upon IFNγ activation, BCL6 is released to permit STAT1 driven transcription. Approaches overexpressing BCL6 or enhancing its binding to DNA (using BTB domain antagonists) sustain BCL6 repression and prevent STAT1-dependent gene expression (**Figure 12**). An increased understanding of the graft-intrinsic processes contributing to rejection would provide the basis for novel tactics to dampen local alloimmune-mediated damage and preserve long-term graft function. Given that endothelial immunogenicity is a central phenomenon in transplant rejection and other vascular diseases, leveraging endogenous BCL6 repressive function may serve to dampen endothelial activation and ameliorate tissue inflammation.

**Figure 12.**
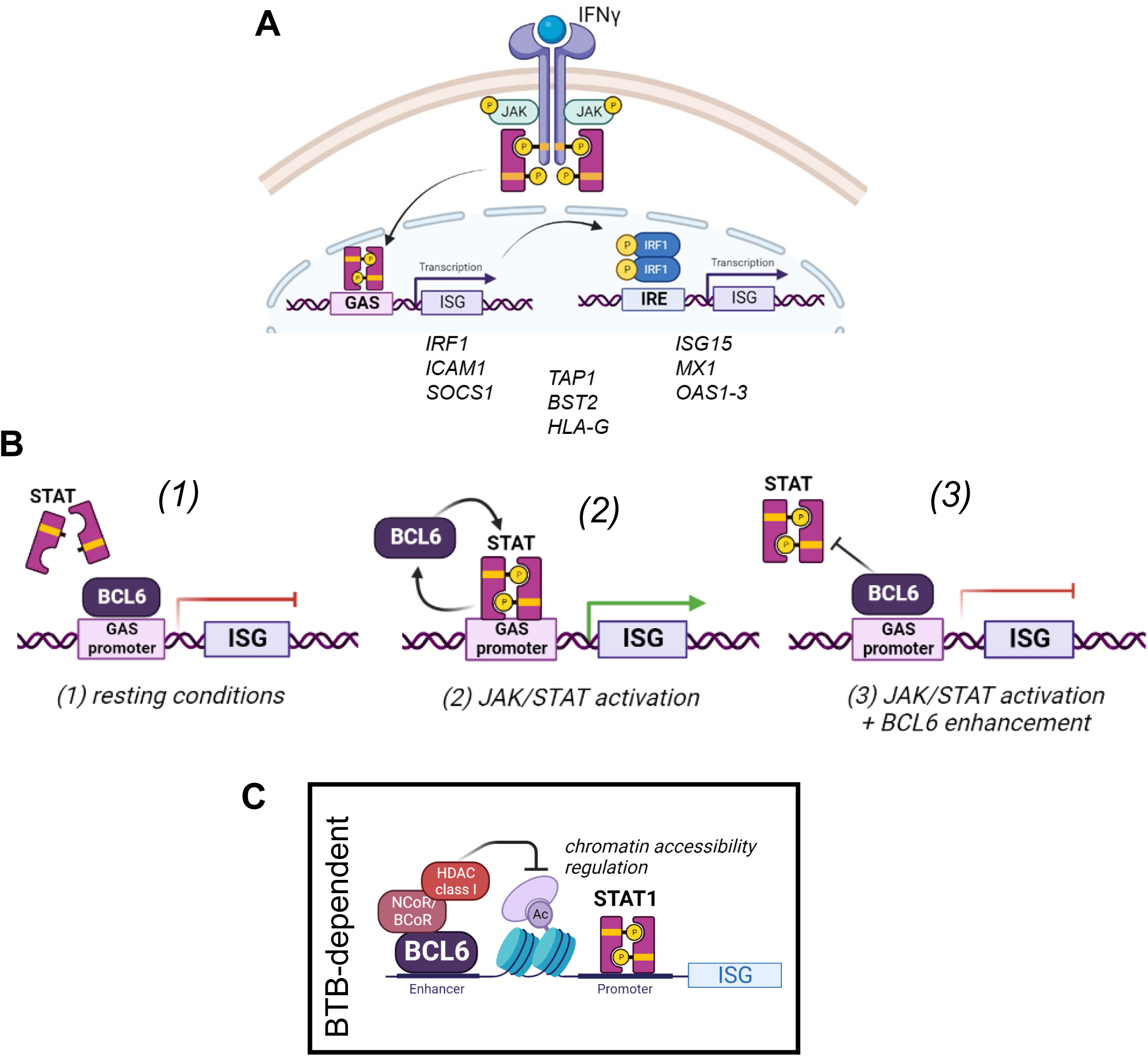
Proposed model of BCL6 regulation of interferon response genes in endothelium. A) Endothelial exposure to IFNγ activates STAT1, which drives expression of GAS-containing target genes including the secondary transcription factor IRF1. IRF1 in turn regulates expression of IRE/ISRE containing genes. B) We propose that BCL6 binds directly to STAT recognition sequences (1), cycling off when STAT signaling is activated to release its repression (2). Here, depleting BCL6 would augment target gene expression by STAT, but in a BTB-independent manner, while overexpression or stabilization of DNA binding would suppress gene expression (3). C) Previous reports show that BCL6 regulates chromatin accessibility through BTB-dependent interaction with histone deacetylases. Here, blocking the BTB domain or depleting BCL6 would be expected to augment target gene expression.

## Acknowledgements

The authors would like to extend their thanks and acknowledge the technical contributions from the following individuals: the UCLA Immune Assessment Core, including E. Cho, M. Rossetti, G. Sunga, and M. Cappelletti, for technical assistance in performing the Luminex assays and development of flow cytometry panels; the UCLA Center for Systems Biomedicine, particularly to E. Faure for performance of Nanostring assays; to the UCLA Translational Pathology Core Laboratory, particularly to Y. Li for immunofluorescence staining of human tissue; the UCLA Technology Center for Genomics and Bioinformatics, especially to X. Li for RNA-Sequencing; and to L. Cerchietti, A. Melnick and M. Koegl for conceptual discussions on BCL6 inhibitors and mechanism of action.

---

**Supplemental Figure 1.**
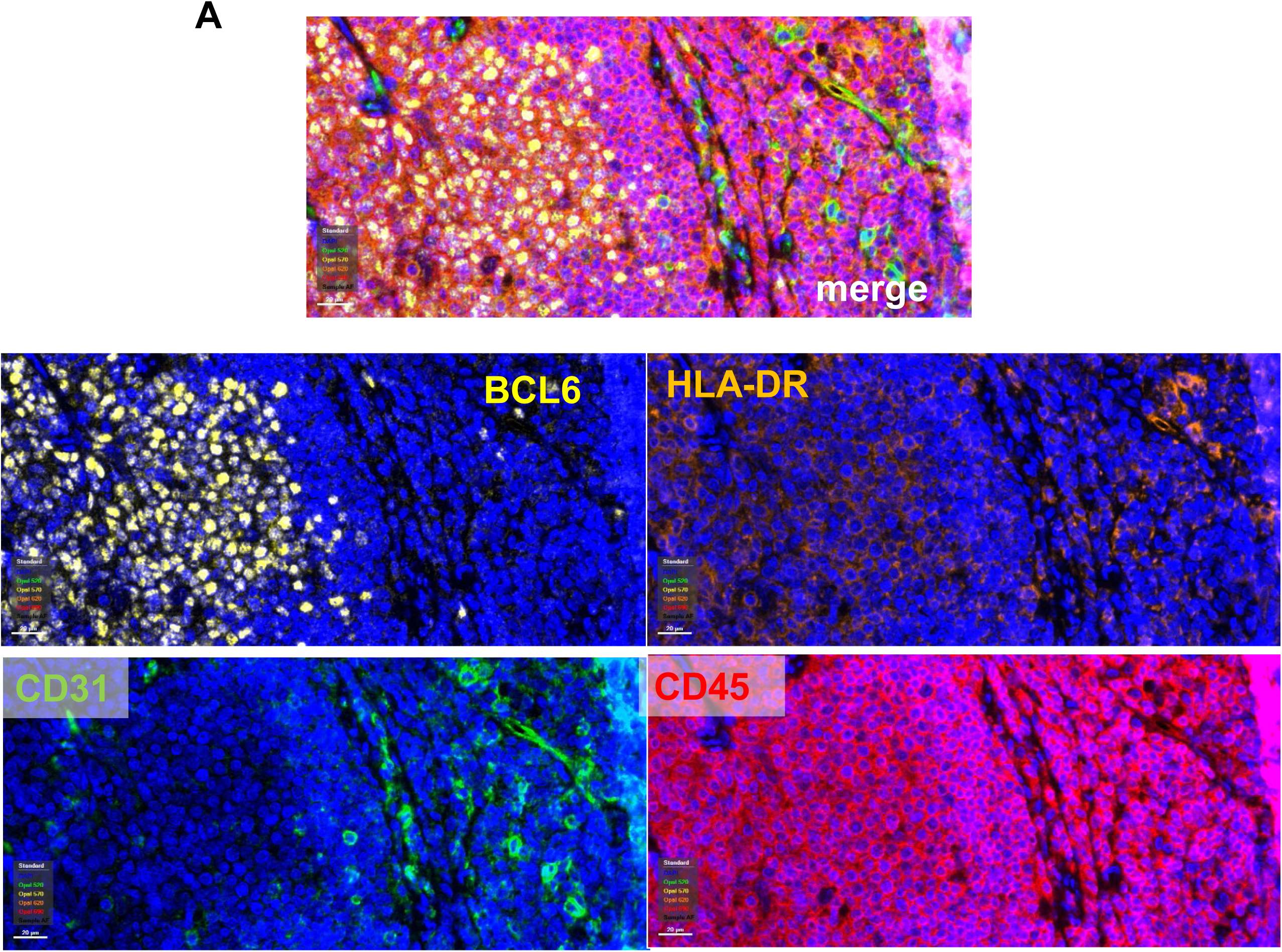
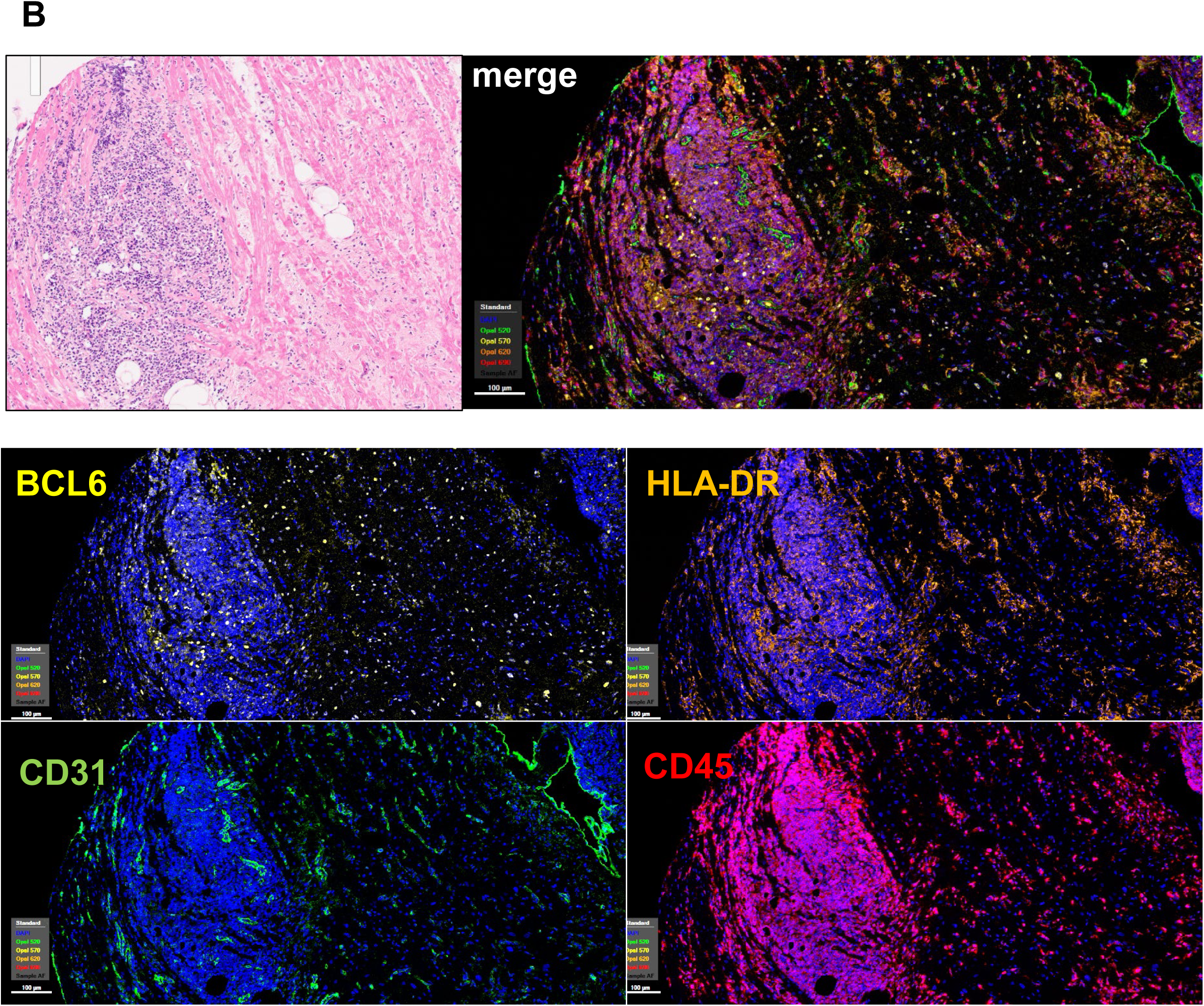
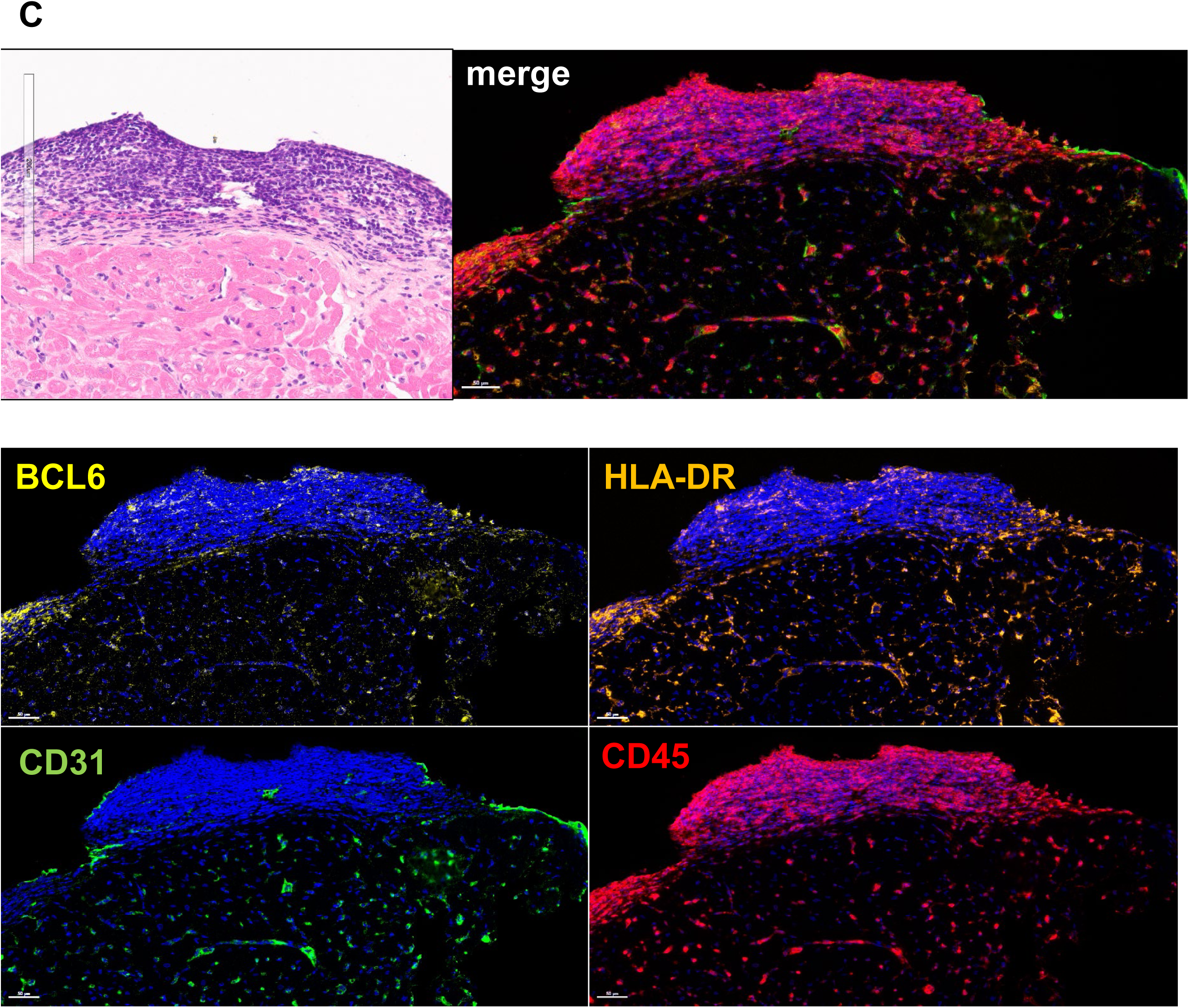
Specificity of antibody staining for BCL6 and HLA-DR in human tissue. A) Human tonsil was tested as a positive control for DAPI, CD31, CD45, BCL6 and HLA-DR staining. B) Exemplary 10X images of a CD19+ Quilty B lesion within a rejecting human cardiac allograft, with numerous BCL6+ cells within the CD45+ lesion. C) Exemplary 20X images of a Quilty A lesion within the endocardium of a rejecting human cardiac allograft, with scant BCL6+ cells within the CD45+ lesion. Top left image shows H&E staining. Top left image is merged staining.

**Supplemental Figure 2.**
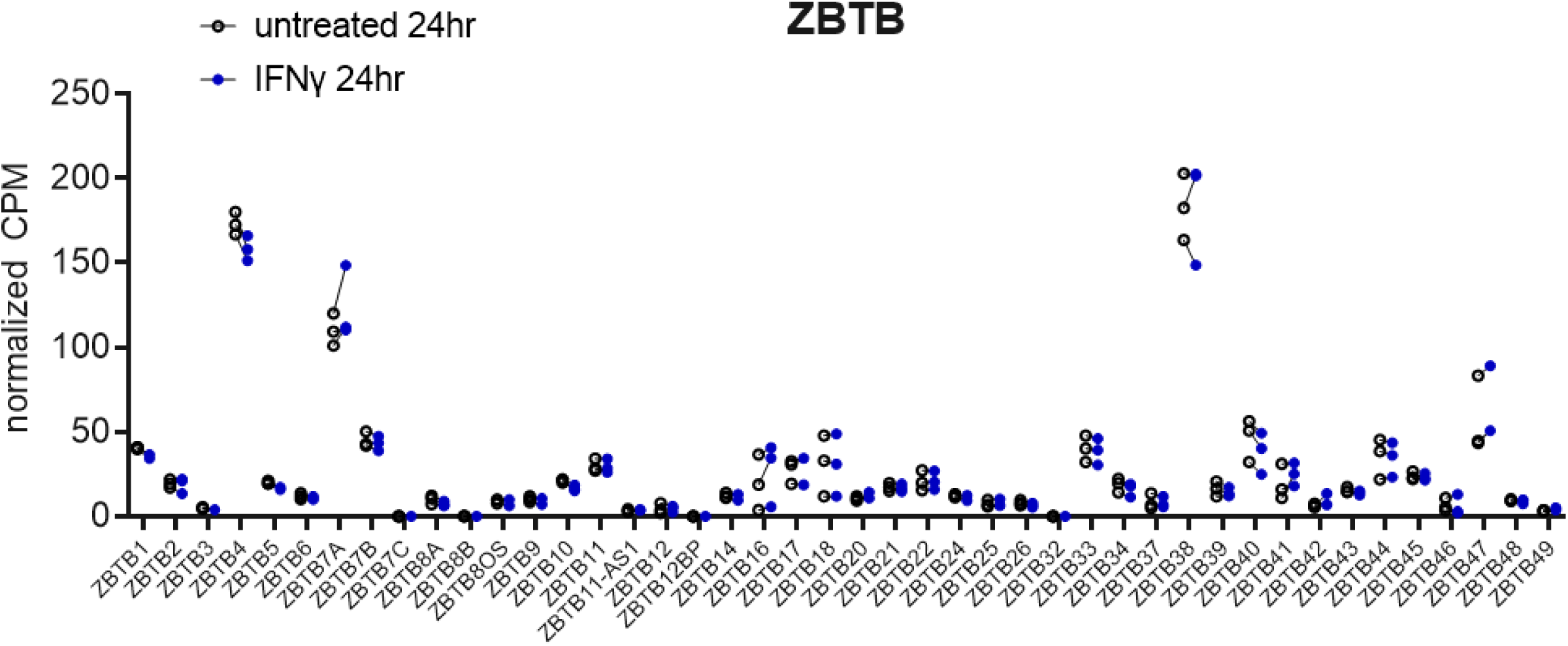
IFNγ had no effect on expression of other ZBTB genes in endothelial cells. Human aortic endothelial cells (n=3 biological replicates) were treated with IFNγ for 24hr and gene expression was measured by RNA-Seq. Normalized CPM for *ZBTB* gene expression is graphed.

**Supplemental Figure 3.**
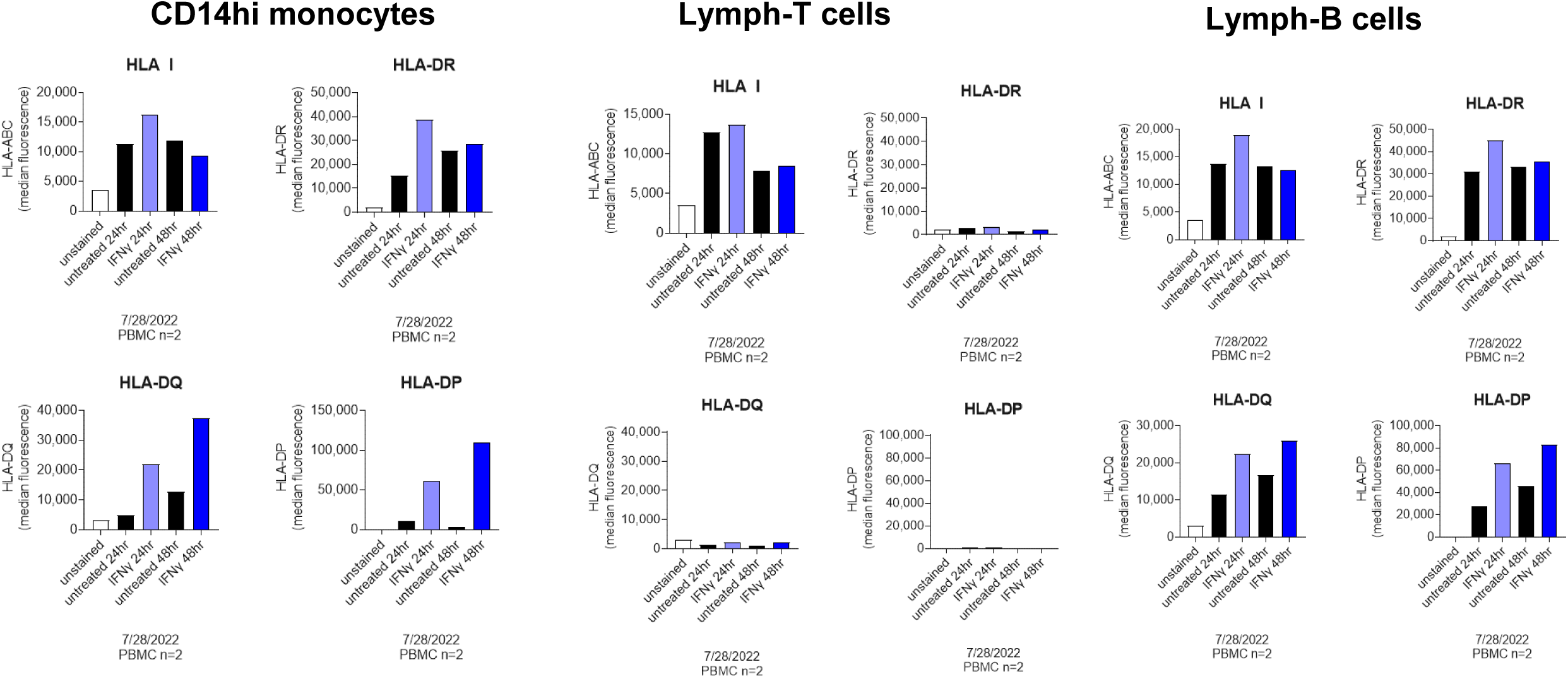
Tempo of HLA class II expression on leukocytes. Peripheral blood mononuclear cells (n=3 donors) were treated with IFNγ for 24hr and 48hr. Cells were then stained for CD14, CD3, CD19, and HLA-DR, -DQ, and –DP by flow cytometry. CD14+ monocytes, CD3+ T cells, and CD19+ B cells were gated in FlowJo, and the median fluorescence of HLA class II was determined for each cell type.

**Supplemental Figure 4.**
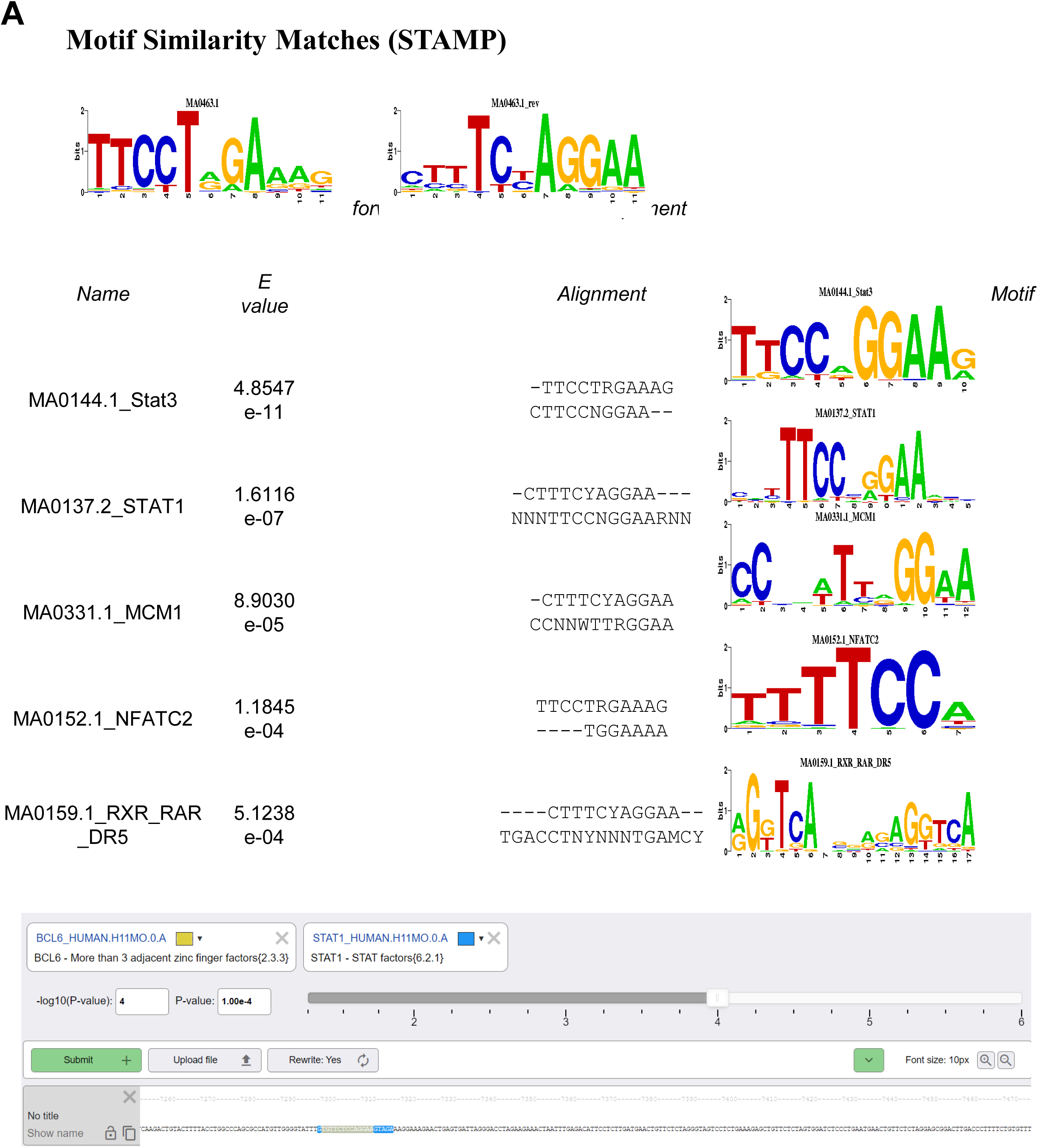
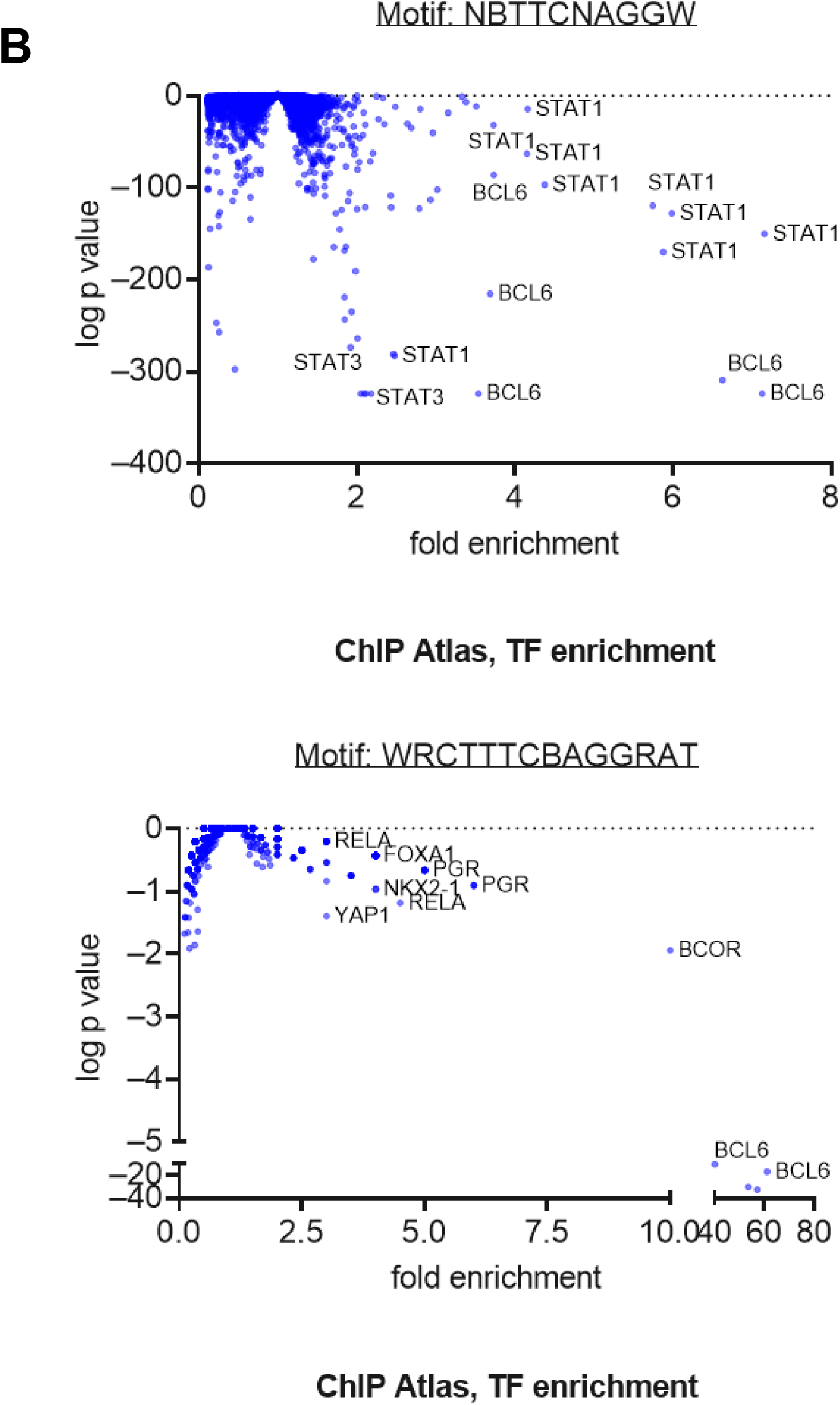
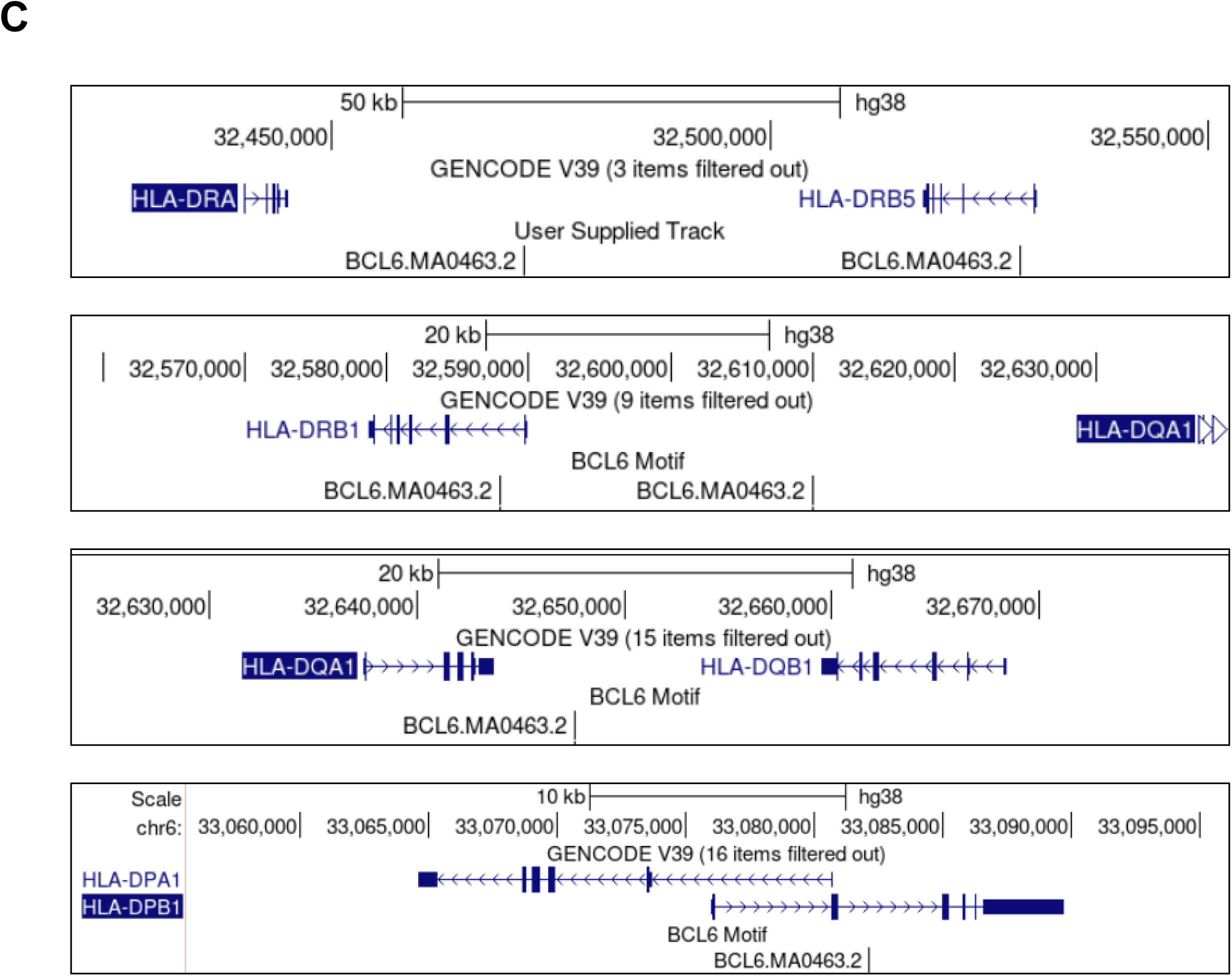
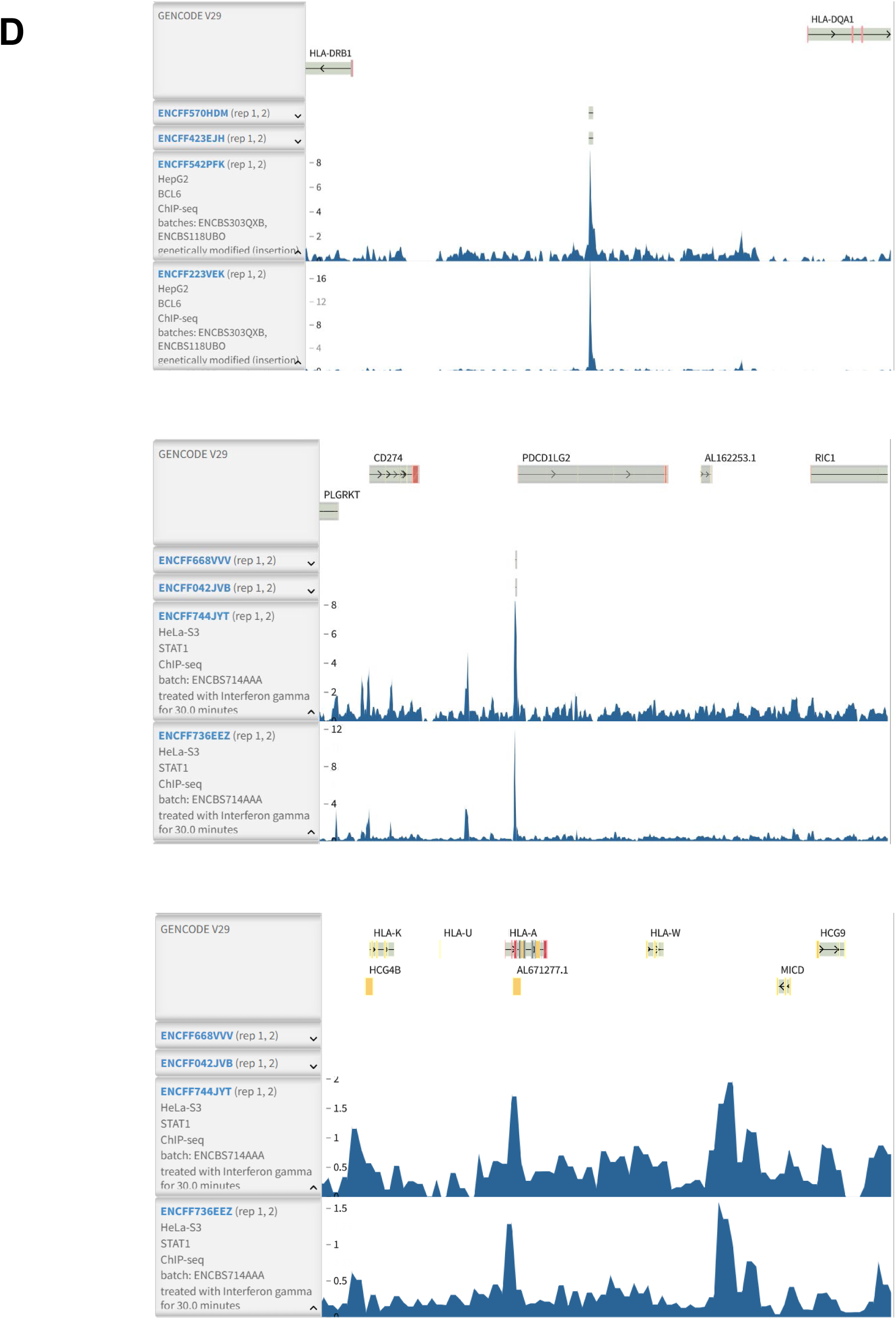
Similarity analysis of the BCL6 consensus motif with other transcription factor recognition sequences. A) The BCL6 consensus motif was analyzed for similarity with other transcription factor binding sites using STAMP. Bottom: the BCL6 and STAT1 consensus sequences were aligned on the human genome and highlighted using molotool. **Comparison of TF enrichment at two BCL6 de novo motifs.** B) Using the unbiased ChIP Atlas Enrichment Analysis, two degenerate consensus motifs for BCL6 were queried against hundreds of public ChIP-Seq datasets for transcription factors with enriched binding to each DNA sequence. The fold enrichment of all transcription factors in the ChIP Atlas database is plotted against the log p value, with annotations for the most highly enriched transcription factors binding to each de novo predicted BCL6 motif. **BCL6 DNA binding motifs are found within HLA class II genes**. C) UCSC Genome Browser view of the BCL6 motif mapped to the human genome (using http://bardet.u-strasbg.fr/tfmotifview/?results=ezFBWpD8uTbPSB), focused on the HLA class II region. **BCL6 DNA binding peaks are found near or at IFN response genes.** D) ENCODE data in both HepG2 and K562 show significant BCL6 binding peaks between HLA-DRB1 and –DQA1 (top), at the TSS of PDCD1LG2 (middle), but not in any classical HLA class I genes (bottom).

**Supplemental Figure 5.**
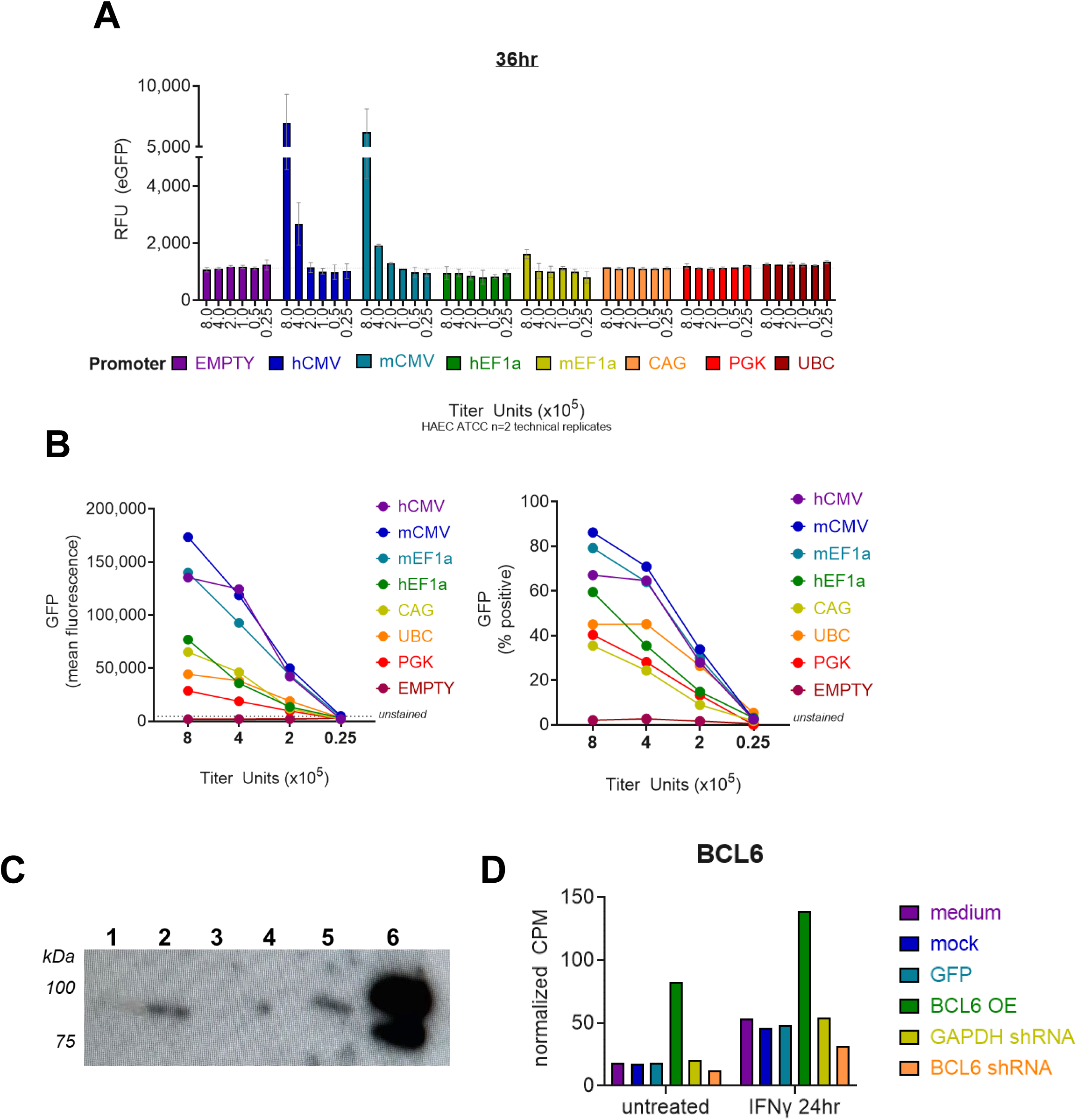
Optimization of lentiviral expression of BCL6 and shRNA. A-B) Promoter activity was screened. Primary human aortic endothelial cells (n=2) were transduced with increasing MOI of lentiviral vectors with GFP driven by a variety of promoters. GFP expression was measured on a Cytation5 fluorometric plate reader 36hr after infection (A) and by flow cytometry 7 days after infection (B). C) Western blot for BCL6 protein expression in transduced TeloHAEC. Lane 1-mock; 2-GAPDH shRNA; 3-BCL6 shRNA; 4-empty; 5-GFP overexpression; 6-BCL6-GFP overexpression. D) Normalized transcript counts per million of *BCL6* in transduced TeloHAEC.

**Supplemental Figure 9.**
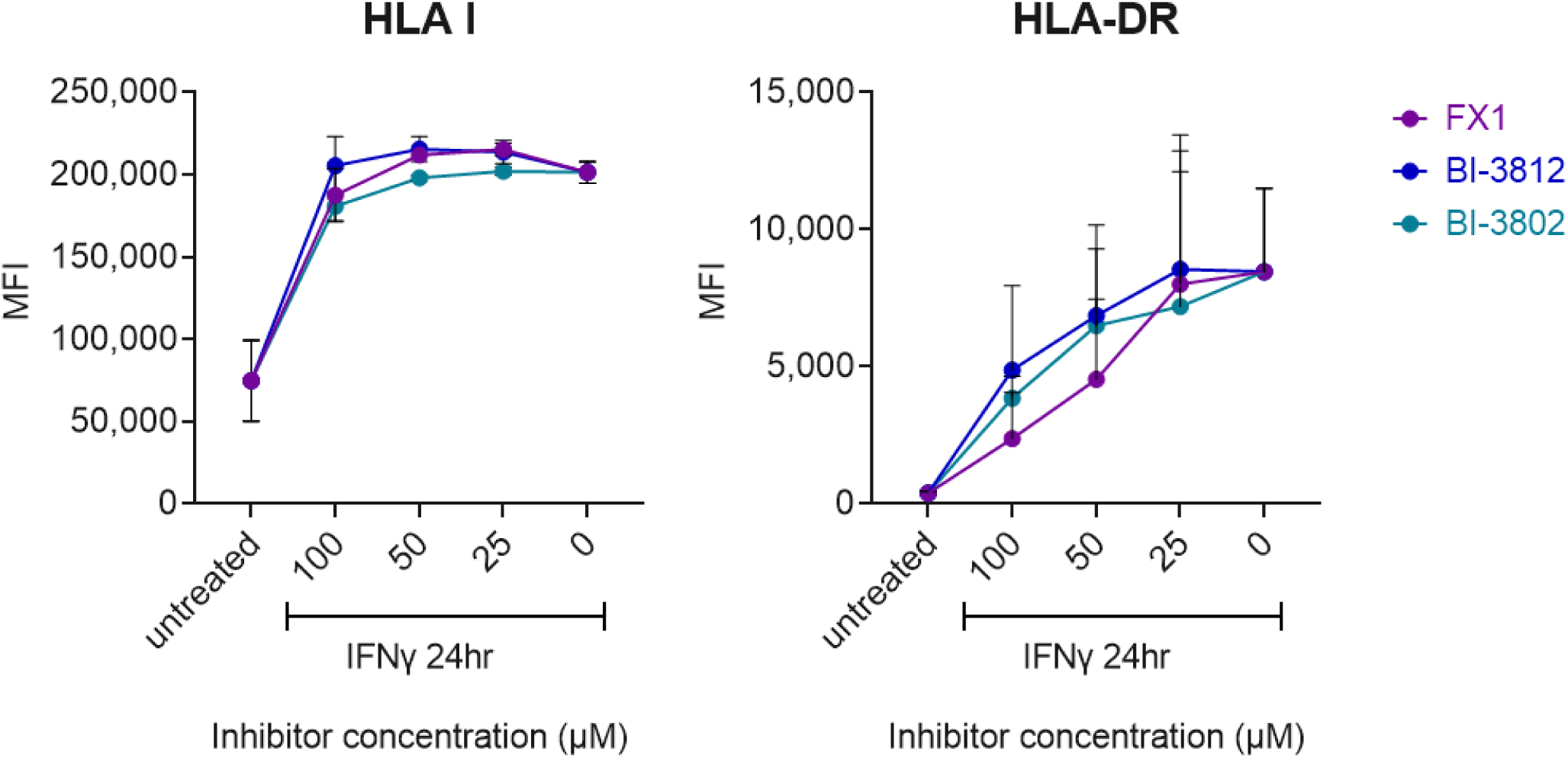
Dose response of BCL6 inhibitors on HLA class I and HLA class II expression. Primary human aortic endothelial cells (n=3 biological replicates) were pre-treated with FX1, BI-3802 or BI-3812 (25-100μM) followed by IFNγ for 24hr. Cell surface expression of HLA-ABC and HLA-DR proteins was measured by flow cytometry.

**Supplemental Figure 10.**
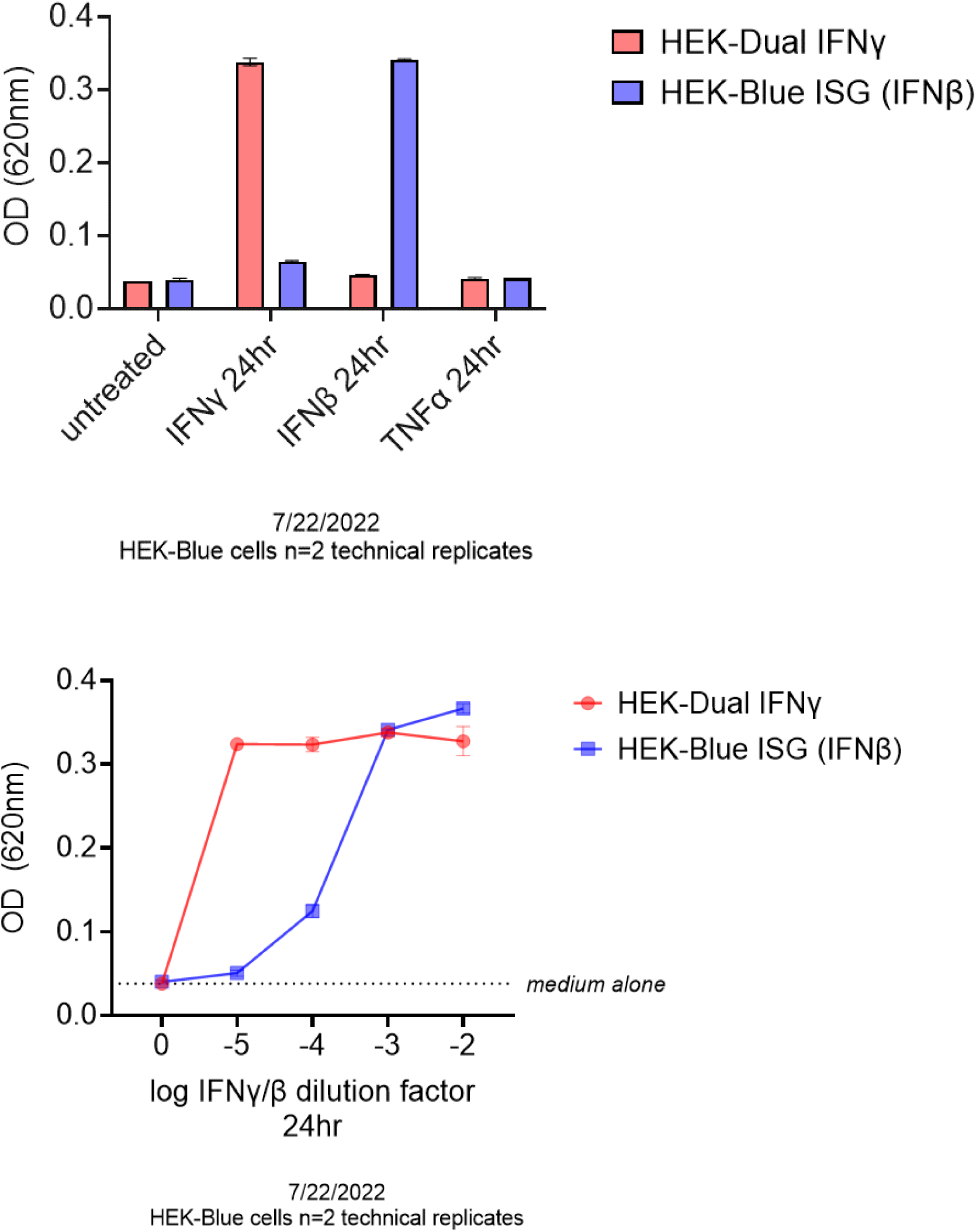
HEK-Blue validation. HEK Blue reporter cells were treated with IFNγ, IFNβ, or TNFα as indicated for 24hr. Transcriptional activity of STAT1 (red) or IRF/ISRE (blue) was tested by SEAP assay. Results show representative technical duplicate values, from 3 independent experiments.

